# Stress-Induced Early Hatching Alters Growth Trajectory and Experience-Dependent Transcriptome in Red-Eyed Treefrog (*Agalychnis callidryas*) Tadpoles

**DOI:** 10.64898/2026.11.15.766833

**Authors:** S.B. Seraphin, K.L. Gendreau, A. Hannifin, S. Dadina, E. Beiga, M. Ryan, J. Altman, R. Rana, S. Phoe, N.L. Vega, A.J. Jacques, M.B. Igayemi, R. Márquez, M. Horb

## Abstract

The ability to grow fast and develop frugally in the face of hazards is a common feature of plants and animals. Life History Theory predicts that trade-offs between immediate survival or reproduction and future longevity may occur under threatening conditions. Embryos of the Neotropical red-eyed treefrog (*Agalychnis callidryas*) hatch prematurely (escape hatching) in response to ecological stressors such as flooding-induced hypoxia, desiccation, and egg-predator attack, making this species an ecologically grounded model for the evolutionary-developmental biology of early life stress (ELS). To assess the physiological correlates of ELS in *A. callidryas*, we bred nine captive adults in a rain chamber and divided ten resulting clutches into sibling-matched Early-Hatch (EH, N = 80; hatching induced by mechanical stimulation upon appearance of the keratinized beak) and Late-Hatch (LH, N = 99; undisturbed) groups. Tadpoles were harvested, measured, and weighed for cross-sectional growth analyses, with subsets used for morphometric and transcriptomic study.

Cross-sectional mass–age trajectories differed between treatments: fitted EH mass rose early and peaked near 12 days post-hatching, whereas fitted LH mass increased throughout the shared 17-day observation interval. Derivative analyses consistently supported a later LH mass–age slope advantage, while evidence for an early EH advantage and greater EH deceleration depended on spline flexibility. Age- and family-adjusted morphometrics (32 EH, 35 LH) revealed treatment differences in overall size but not size-independent shape. Gene expression analyses of three sibling-matched pairs identified 258 differentially expressed transcripts suggestive of a delay in adaptive immune system development and increased oxygen and antioxidant demands in EH. Threatened *A. callidryas* embryos do not merely hatch early: their premature entry into tadpole life initiates a developmental program of early compensatory growth followed by pronounced deceleration, accompanied by transcriptional differences consistent with life-history trade-offs between immediate survival and later development.

## INTRODUCTION

Stress is universal. It requires alterations to the organism’s physiology in support of immediate demands for environmental adjustment. Life History Theory (LHT) predicts certain tradeoffs, between immediate survival or reproduction, and future longevity, may ensue under threatening conditions (Stearns, 1998, 1976). Proximate risk and ultimate consequences must be weighed against necessary allostatic adjustments (Nettle et al., 2013; Nettle and Frankenhuis, 2020). In keeping with reaction norms, organisms may accomplish allostasis through behavioral, anatomical, or phenological changes both visible and invisible to the naked eye (Cooper et al., 2025).

Behavioral adjustments to stress can include altered social competition, cooperation, dietary flexibility, or niche creation. For example, intense social conflict may lead to the fissioning of social groups through separate foraging or migration (Chapman et al., 1995; Lerch et al., 2021; O’Bryan et al., 2020; Wakeford et al., 2018). Similarly, physiological adjustments to stress can be achieved by changing resource allocation between growth, maintenance, and reproduction. In a survey of the literature, we observed that many species of animals and plants alter the timing and tempo of development, maturation, or reproduction, when confronted by acutely distressing environmental conditions. Long before the appearance of morphological and physiological alterations associated with variable environmental conditions, there may be stress-induced changes to when specific developmental events occur (timing) or how quickly development progresses (tempo).

For timing, both early and delayed onset of developmental maturation are documented in natural and experimental settings. Precocious development seems to be linked with acute environmental uncertainty or instability. For instance, early developmental timing has been observed in golden hamsters exposed to social aggression (Kollack-Walker et al., 1997), rats given insufficient bedding material but fed a high calorie diet (Strzelewicz et al., 2019), heifers weaned early but fed a high-corn diet (Gasser et al., 2006), and sea urchins experiencing crowding stress (Jian et al., 2021). By contrast, environmental instability resulting in increased mortality signals the need to capitalize on immediate opportunities and hasten reproduction (Yang et al., 2022). Despite breeding continuously and persisting throughout the year in tropical regions, annual plants like the Petunia (*Petunia hybrida*) and ephemeral insects like the Green Peach Aphid (*Myzus persicae*) will complete their entire life cycle within one season in less favorable climates (Blackman, 1974; Sink, 1984; van der Krol and Nam-Hai, 1993).

Conversely, poor resource conditions may signal the need to reduce growth and delay reproduction. Thus, the European perch (*Perca fluviatilis*) and Bluegill (*Lepomis macrochirus*) can delay maturation in nutrient-poor, oligotrophic lakes (Rolls et al., 2017). While malnutrition is a common environmental cause, seasonal cues associated with diminishing light can delay subsequent puberty in lambs born during the fall (Foster et al., 1986). Additionally, limited opportunities for reproduction due to the scarcity of food or territory, may delay sexual maturation and necessitate raising siblings among cooperatively breeding bird species (Ball and Wade, 2013; Brown, 2014). By necessitating different morphological and physiological adjustments, different early environments yield divergent morphometric responses (Vehanen and Huusko, 2011). In nature, a multitude of complex environmental variables can coalesce and intensify phenotypic responses required. Hence, morphological and physiological adjustments to temperature variability over a 12-year period were more pronounced in wild sea trout (*Salmo trutta L.*) compared to hatchery-reared sea-run brown trout (Lejk et al., 2021).

Regarding tempo, slow and fast rates of developmental maturation have been observed. Notably fast, Killifish (*Nothobranchius furzeri*) can enter a developmentally quiescent state (diapause) or accelerate their developmental trajectory to suit environmental conditions of drought or more favorable ephemeral pools (Reichard and Polačik, 2019; Vrtílek et al., 2018). Here, early hatching, fast maturation and spawning are rooted in mechanisms of embryo morphogenesis (altered cell cycling during cleavage) in a manner that is both sensitive to environmental stresses and intra-population variation (Dolfi et al., 2019, 2014; Vrtílek et al., 2018). For examples of slower tempo, the offspring of female mice stressed immediately before gestation had slower physical maturation in characteristics such as teeth, weight, length, dexterity and certain reflexes (Meek et al., 2000). Snake eggs incubated at low temperatures take longer to hatch (Burger, 1991). Both delayed growth and menarche have been observed in girls with high urinary cortisol (Shi et al., 2011).The emotional stress of social subordinance has also been linked to slower tempo of sexual maturation in female rhesus monkeys (Wilson et al., 2013).

By adjusting developmental tempo and timing, organisms facing threats from predation, social competition, or environmental hazards can alter the timeline for crucial ontogenetic niche shifts between life stages and habitats to improve fitness. For instance, many Old and New World tropical amphibian species will hatch early under conditions of parental neglect, predator threat, or ecological distress. In multiple frog lineages that have independently evolved terrestrial eggs and retained aquatic tadpoles, including phylomedusine treefrogs (Gomez-Mestre et al., 2013) and glassfrogs (Delia et al., 2019) embryos hatch and enter the aquatic environment early upon exposure to ecological (e.g., predation, flooding) or social (parental neglect) stressors.

As a broad phenomenon, similar environmentally sensitive shifts in hatching timing have been documented in fishes, reptiles, and invertebrates (Du and Shine, 2022; Warkentin, 2011).

As a response to early life threats, post-hatching compensatory growth in frogs and fishes represents an immediate predation-risk trade-off, which may be accompanied by a loss of physical reserve for subsequent maturation (e.g., metamorphosis) or longevity –costs resulting in a shortened lifespan. The “cognitive ecological” processes through which developing organisms integrate environmental cues, assess risk, and make plastic niche-shifting decisions (Touchon et al., 2013; Warkentin, 2011) may be shaped by gene–environment interactions that influence developmental stability and morphology (Gomez-Mestre et al., 2010; Touchon and Wojdak, 2014). Fluctuating asymmetry can serve as one phenotypic indicator of variation in developmental buffering, although its relationship to environmental stress and fitness is context- and trait-dependent (Klingenberg, 2019; Leamy and Klingenberg, 2005).

While escape hatching is not a result or example of “developmental acceleration” per se, it represents a truncation of the embryonic stage resulting in early onset of the tadpole stage. Precocious puberty, early senescence, and elevated morbidity are demonstrated consequences of early life stress in humans. Considering the potential for shared drivers of accelerated developmental timing in humans with escape hatching and comparable advancement of later developmental stages in other species, it may be that early distress disrupts typical development by altering the expression of genes involved in neurobiology, immunity, and ageing. A tractable laboratory animal model could greatly illuminate the physiological mechanisms underlying environmental stress induced changes in developmental timing.

Red-eyed treefrogs are a good candidate species because they are neither endangered nor threatened, and captive-bred animals can generate large quantities of eggs under proper conditions. Furthermore, their developmental timeline from embryo to froglet is relatively short (a few months) considering their moderately long (10 years) lifespans (Touchon et al., 2013; Vitt and Caldwell, 2013). Finally, as an animal model for early life stress (ELS), *A. callidryas* fulfills the important criteria such as the phenotypic response of developmental acceleration (face validity). It also has a potential for shared underlying phylogenetic, ontogenetic, proximate, or ultimate determinants (construct validity), and typical outcomes of tradeoffs in immunity (predictive validity). We hypothesized that stress-induced early hatching would lead to early compensatory growth that is accompanied by altered morphology and gene expression, consistent with adaptive trade-offs.

## MATERIALS & METHODS

### I. Research Subjects

Research animals included 9 captive-bred adult, red-eyed treefrogs (*Agalychnis callidryas*) (Josh’s Frogs, LLC.: 3 males, 6 females), 182 unique laboratory-bred tadpoles from 10 clutches assigned to Early-Hatched (EH, N = 80) and Late-Hatched (LH, N = 99) treatments or control (N = 3) groups. *A. callidryas* are native to forests in Southern Mexico, Central America, and northwestern South America. They inhabit warm, humid lowland forests, typically near ponds, swamps, or other still water bodies used for reproduction (IUCN Amphibian Specialist Group, 2016; Vitt and Caldwell, 2013). Although they retain the ability swim, adults are generally arboreal, nocturnally active, often sleep on the undersides of leaves during the day, and require humid, well-ventilated conditions; eggs are deposited on vegetation over water and the aquatic larvae develop as tadpoles (Vitt and Caldwell, 2013; Warkentin, 1995). They are currently listed as a species of least concern, with regard to conservation (IUCN Amphibian Specialist Group, 2016).

In captivity, a daytime thermal gradient of approximately 75–85°F (24–29°C), a modest nighttime decrease to 68–75°F (20–24°C), and relative humidity of about 70–75% during the day with higher nighttime humidity are appropriate husbandry targets (Exo Terra, 2023). Reported longevity varies; wild individuals are often estimated to live about five years, whereas captive animals may live eight years or longer under suitable care.

#### Animal Husbandry

Research was conducted at two sites: 1) Laboratory of Evolutionary Neuroscience, Trinity College (Hartford, CT USA) and 2) Laboratory of Marko Horb, Marine Biological Laboratory (MBL, Woods Hole, MA USA). All procedures were approved by their respective Institutional Animal Care and Use Committee (Trinity IACUC Protocol #: 67-22 and 67-25; MBL IACUC Protocol #: 22-27, 23-32, and 24-34), in keeping with State and Federal guidelines.

##### i. Adult Frog Housing

Red-eyed treefrogs were housed under naturalistic conditions (Touchon and Hughey, 2025), in mixed-sex groups of 2 – 4, within 12x12x18 inch (2926.08 cm^2^) bioactive glass terrariums outfitted with plants, bark for climbing, and a substrate of Sphagnum moss, leaf litter or coconut mulch, as well as beneficial microfauna (New England Herpetoculture, LLC: Springtails (Collembola) and Woodlice (Isopods)). Terrariums were maintained in a colony room with an automatic 12-hour light, 12-hour dark cycle. During the light phase, additional illumination from UVA/UVB bulbs (Zilla®, 13-Watt Fluorescent UVB/UVA Bulb) was used to facilitate photosynthesis of vitamin D3 necessary for calcium absorption and metabolic disease prevention. Programmable misters (Exo Terra Monsoon SOLO II) kept terrarium average minimum humidity to 69.6% (Range: 66.2 - 77.5%) and average maximum humidity to 79% (Range: 71.5 - 81.6%). Dome lamp fixtures fitted with 24-hour warming lights (Zoo Med® 75-Watt Infrared Heat Lamp) maintained terrariums at an average daily temperature of 73.8°F (69.4 - 79.1) while preserving nighttime darkness.

##### ii. Pre-Metamorphic Frog Environments

Tadpoles and metamorphing frogs were maintained, at a density of approximately 2 animals per quart, in 5.21 L aquariums adapted from clear, lidded, and food-grade (BPA Free) plastic storage containers (Elsjoy), into which 1/16” (1.59 mm) holes were manually drilled in each corner, for air circulation. Indian Almond Tree (*Terminalia catappa*) leaves are rich in tannins and flavonoids, which have beneficial effects in the culture of aquatic organisms (Sung and Abol-Munafi, 2020; Wang et al., 2024). For enhanced acidity, and its antifungal and antimicrobial properties, a blackwater medium (mean pH 7.25) was prepared by steeping ∼15 Almond leaves (Josh’s Frogs, LLC) in 5 Gallons (18.93 L) of cold, distilled water overnight. Thereby, water quality was maintained at an average pH of 6.8 (6.5-7.1), ammonia level of 0.4 ppm (0-1), and a nitrate/nitrite level of 0 ppm (0-0) with an average temperature approximately the same as room temperature (73°F).

While tanks containing small tadpoles were aerated with a bubbler (Tetra® Whisper Air Pump) to minimize turbulence, tanks with large tadpoles and metamorphing frogs were mechanically filtered (Tetra® Whisper Internal Power Filter). Tadpoles were provided with sections of 4-inch (10.16 cm) diameter Polyvinyl Chloride (PVC) pipe shelters, laid horizontally, for use as hideouts. Metamorphs were also supplied with rocks or platforms on which to climb above the water surface.

##### iii. Tadpole and Metamorph Care

Each filter-feeding tadpole was supplied with 3.5mL of 2gm/50mL Sera® Micron Nature fry food containing 18% krill (zooplankton) and 51% spirulina (phytoplankton). Upon evidence of bottom feeding, larger tadpoles received high protein pellets (Zoo Med Aquatic Frog & Tadpole Food) or bloodworms (Hikari® Bio-Pure Freeze-Dried Blood Worms). Metamorphing frogs and froglets were fed fruit flies (Wingless *Drosophila melanogaster*, or Flightless *D. hydei*), 1/16-inch crickets (Pinhead Banded Crickets, *Gryllodes sigillatus*)).

##### iv. Juvenile and Adult Frog Care

Adult frogs received ad libitum access to bowls of water that was purified by reverse osmosis or distillation and re-mineralized with a conditioner (Josh’s Frogs, LLC: R/O Rx Distilled Water Remineralizer). Animal care technicians also introduced live insect feed (Josh’s Frogs, LLC: ¾”-1” Banded Crickets (*Gryllodes sigillatus*) or Orange Head Roaches (*Eublaberus posticus*)) to terrariums daily. Insects were dusted with the manufacturer recommended amount of Rep-Cal, Calcium Plus once per week.

### II. Experimental Protocols

#### Frog Breeding, Egg Harvesting and Herpetoculture

##### i. Frog Reproduction

To simulate rain that typically precedes frog mating, we employed an engineered “rain chamber” for mating 9 adult *A. callidryas* (Josh’s Frogs, LLC: 3 males, 6 females). The rain chamber consisted of a 12.4” deep x 24.2” high x 24.2” wide (15,273.5 cm^2^) glass paludarium donated by Dr. Karen Warkentin (KW, Boston University). After adding plastic and live tropical plants for use as substrates, this was fitted with a submersible pump (Exo Terra Repti Flo 200 Circulation Pump) to circulate purified water, from the tank basin and up through a custom-perforated plexiglass lid from KW. A timer was programmed to operate the pump on the following water schedule: short 1–5-minute bursts of rain every 4 hours; long 2-hour “torrential” rain showers at 8:00pm and 12:00am. This continued for 2 - 3 nights, during which males exhibited increased vocalization, displacement aggression, and amplexus with females (Pyburn, 1970). The breeding regimen generated over 200 eggs across three breeding cycles.

##### ii. Egg Collection

Each morning following a night of breeding, the rain chamber was visually inspected for the presence of eggs. With care not to disturb adult frogs, eggs were carefully relocated by either snipping off the leaf on which a clutch was laid or using a silicon spatula to gently remove the egg mass. After collection, eggs that appeared white and without signs of cellular division were presumed unfertilized, carefully removed with tweezers, and discarded. Each clutch was assigned an identification number and censused.

##### iii. Semi-Randomized Clutch Separation and Study Assignment

Using a silicone spatula, each clutch was then divided in half, down the midline, to form sibling-matched “Early-Hatch” (EH) and “Late-Hatch” (LH) treatment groups. A diagram showing clutch division/separation can be found in [Figure 1].Taking care to maintain the original bottom-down orientation, the EH and LH batch from each clutch was then adhered to a 3 X 5” (7.62 X 12.7 cm) food-grade plastic card made from pliable, bacterial-resistant cutting boards. The cards were affixed, using duct-tape, to the inside of different 5.5 Quart (5.21 L) aquariums for separate incubation. To provide a water source for eventual hatchlings, these mini aquariums were filled with 3” (7.62 cm) of almond leaf water.

**Figure 1:**
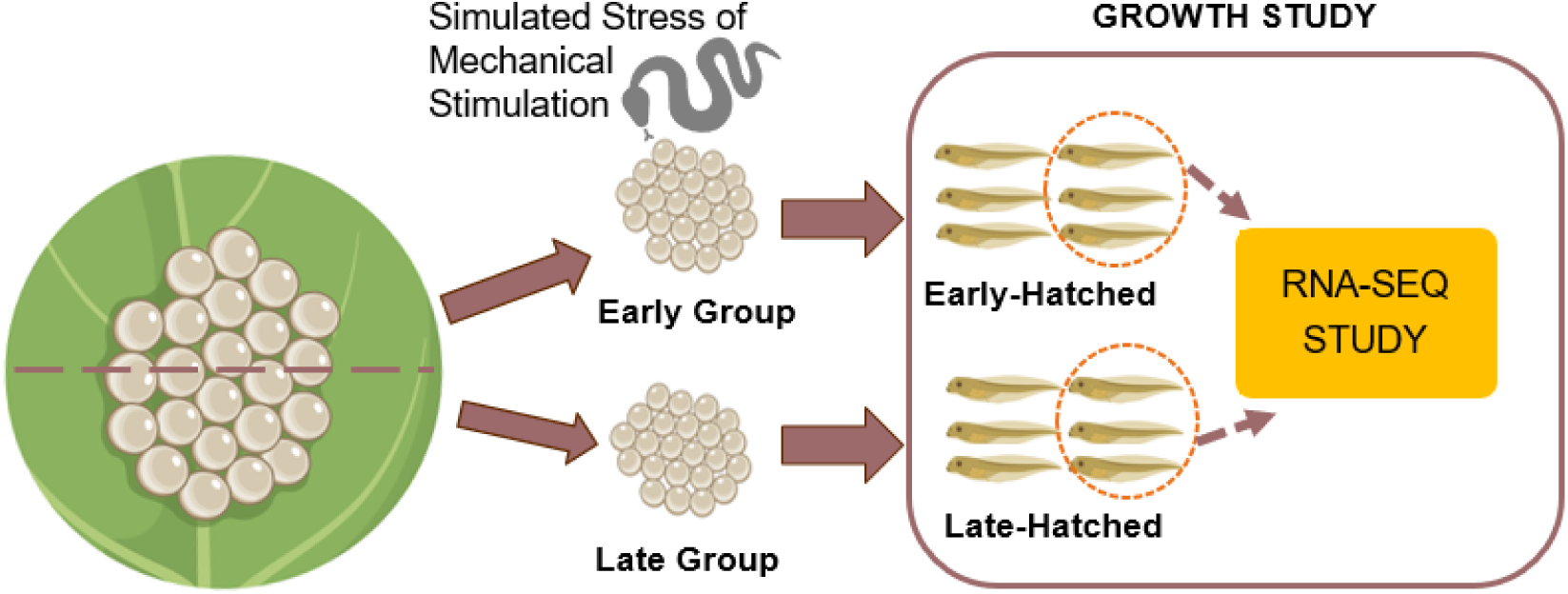
Differential Treatment and Tissue Sampling Protocol. Diagram depicts the hatching treatments and tissue-sampling protocol. The full EH/LH sample comprised 80 EH and 99 LH animals; mass analyses used the 75 EH and 98 LH animals with available harvesting mass. LH were not disturbed and spontaneously hatched. The escape response was induced in EH, by stroking eggs with gloved hands for 15 sec intervals to simulate the mechanical stimulation of a snake predator (stressor). The transcriptome response to stress-induced or undisturbed (RNAseq study) was compared between three EH animals and their matched LH siblings.

##### iv. Embryo Culturing

To create a humidified chamber for embryo culture, an 18 Qt (17.03 L) capacity, 17.2” X 22.8” X 12.5” (43.688 cm X 57.912 cm X 31.75 cm) plastic storage container (Hefty 18 Qt Hi-Rise Stackable Plastic Storage Bin) was adapted by drilling a 5mm hole into the lid to accommodate the tubing from an automatic fogger (ZooMed Repti Fogger™ Terrarium Humidifier). Up to 4 mini aquariums were placed in each humidified chamber, based on EH and LH treatment conditions. In addition, a few eggs were cultured in a petri dish so that developmental progress could be monitored under a microscope without disturbing the treatment groups. Ensuring that the fogging nozzle pointed away from the eggs, all eggs were maintained at room temperature, under approximately 80% humidity, and monitored for signs of excess moisture (dramatic swelling), insufficient moisture (shrinkage), and rot (whitening, graying, or blackening). On days 1 – 2 post laying (before the development of mechanosensory abilities in the embryos or when early hatching can occur), spoiled eggs were carefully removed using direct suction from a transfer pipet. After this, lab personnel took extra precautions to not jostle, shake, or otherwise roughly handle the humidifying chambers at any time during the incubation period.

##### v. Developmental Staging

Using the petri specimens, embryos were periodically observed under microscope (ZEISS SteREO Discovery.V12), beginning at the 16-cell stage onward. Using two developmental atlas systems: Gosner Stages (GS; Gosner, 1960) and Warkentin Stages (WS; Warkentin, 2017), they were monitored for signs of development progression through developmental stages. This included the appearance of the neural tube (GS15), heartbeat (GS15), the proliferation of external gills (WS20-22), and the appearance of melanophores (WS25) and parallel eyes (WS29) (Gosner, 1960; Pyburn, 1970; Warkentin et al., 2017). Although mechanosensory-cued early hatching can occur as soon as GS23 (Warkentin, 2017), we used the presence of keratinized beaks at WS30 to mark the experimentally induced early hatching.

##### vi. Embryo Hatching Procedures

In *A. callidryas* “Early Hatching” (EH, N=80) was experimentally induced by mechanical stimulation circa Gosner stage 30, where keratinized beaks are visible and mechanosensory abilities are developed. This was accomplished by having a researcher, wearing gloves, gently rub the surface of the egg mass, passing back and forth at 15 second intervals followed by 15 seconds of rest (no stimulation). The start and stop of protocol was intended to artificially simulate the vibration rhythm of predator attack, which typically includes a period of pause for swallowing (Jung et al., 2022).

Conversely, the “Late Hatching” (LH, N=99) condition for in *A. callidryas* required no intervention. These embryos were left to hatch on their own, without disturbance or interference. The humidified chambers were only observed in the morning and afternoon so that the date of hatching could be noted for comparison with their EH counterparts. A diagram depicting this different treatment can be found in Figure 1. LH typically hatched within 24 hours of their siblings. On a few occasions, where there was partial-clutch hatching of LH tadpoles an “average” hatching-date was assigned to that clutch, and they were excluded from this study. Three additional 5-day embryo control specimens (GS 30-33 range) were decapsulated on ice. These specimens were retained as descriptive pre-hatching mass references and were excluded from all EH– LH inferential analyses. They were not used for anchoring the primary GAM predictions.

### III. Tissue Harvesting, Measurement, and Preservation

At regular intervals post hatching, sibling-pairs of tadpoles were randomly selected, from EH and LH treatment groups. To build a developmental timeline, effort was made to harvest 1-2 animals from each condition daily. Upon collection, they were measured and, depending on downstream use, preserved for genetic or immunohistochemistry procedures. Animals were euthanized by immersion in 0.1% pH 7.0 MS222 (Sigma-Aldrich, Inc.: Tricaine Methane Sulfonate) for 15 minutes, rinsed in purified water, and then weighed. Since each individual tadpole was not followed longitudinally, this dataset is cross-sectional. A single wet body mass measurement was obtained at harvesting, for each tadpole, so Weight x Age and Gene-Expression x Age relationships could be studied between EH and LH. Specimens from EH or LH sibling-pairs were then preserved for future genetic and histological studies as follows: 1) one tadpole from each EH or LH pair was preserved in 400uL Invitrogen™ TRIzol™ Reagent (Thermo Fisher Scientific, Inc.) and stored at -80°C until RNA extraction. 2) the second was preserved in MEMFA (1M MOPS, 20mM EGTA, 10mM MgSO4, 38% Formaldehyde) for histology.

***i. Age Post-Laying Sampling Time* –** Across 179 unique Early-Hatched (EH; n = 80) and Late-Hatched (LH; n = 99) tadpoles, mean age post-laying (APL) at sampling was 17.23 days (SD = 4.38; range = 8–25 days). EH tadpoles were sampled at a mean APL of 16.93 days (SD = 4.91; range = 8–25), whereas LH tadpoles were sampled at 17.47 days (SD = 3.90; range = 10–25). APL variance was greater in EH than LH tadpoles (variance-ratio F(79, 98) = 1.59, p = .029; median-centered Levene W = 4.50, p = .035), but mean APL did not differ between treatments, Welch t(148.75) = −0.81, p = .417.

***ii. Age Post-Hatching Sampling Time* –** Across the same 179 tadpoles, mean age post-hatching (APH) at sampling was 9.77 days (SD = 5.26; range = 0–20 days). EH tadpoles were sampled at a mean APH of 10.58 days (SD = 5.83; range = 0–20), whereas LH tadpoles were sampled at 9.12 days (SD = 4.69; range = 0–17). APH variance was greater in EH than LH tadpoles (variance-ratio F(79, 98) = 1.55, p = .041; median-centered Levene W = 4.68, p = .032). The difference in mean APH did not reach statistical significance, Welch t(150.10) = 1.81, p = .073.

***iii. Data Quality Control and Sensitivity Analysis* –** Inspection of the harvesting records identified a potentially anomalous collection batch comprised by five late-hatched specimens (14A.7, 14A.8, 14A.9, 16AB.7, and 16AB.8). All five were laid on June 2, 2023, hatched on June 9, and harvested together on June 21 at 12 days post-hatching, yet their recorded masses ranged from 21 to 368 mg. This dispersion was unusually large relative to other LH specimens sampled at the same post-hatching age, and the five records also lacked the accompanying morphometric measurements available for most specimens. Because there was no definitive evidence that the recorded masses were erroneous, all observations were retained in the primary analyses. As a quality-control sensitivity analysis, all principal growth models were repeated after excluding only these five specimens to determine whether treatment effects depended on this potentially anomalous collection batch.

***iv. RNAseq Subjects Sampling –*** A representative subset of six tadpoles were selected for RNA sequencing. These were three family-matched differently treated sibling pairs. All subjects originated from clutches laid on June 14, 2022; within each family, one tadpole from the EH clutch and one from the LH clutch were sampled. Harvest ages spanned 0–12 days post-hatching and 8–22 days post-laying, providing paired family comparisons across early development, although paired siblings were not harvested at identical ages [Figure 2]. EH siblings were heavier on average (62.3 mg) than LH siblings (36.7 mg), with a mean paired difference of 25.7 mg; but this estimate is highly imprecise with only three pairs. The paired comparison was not statistically significant: paired t(2) = 1.19, p = .357. The exact Wilcoxon signed-rank test was also non-significant (p = .50) [Figure 3]. Although under-powered statistically, this analysis supports the assumption of physical similarity between EH/LH sibling pairs.

**Figure 2:**
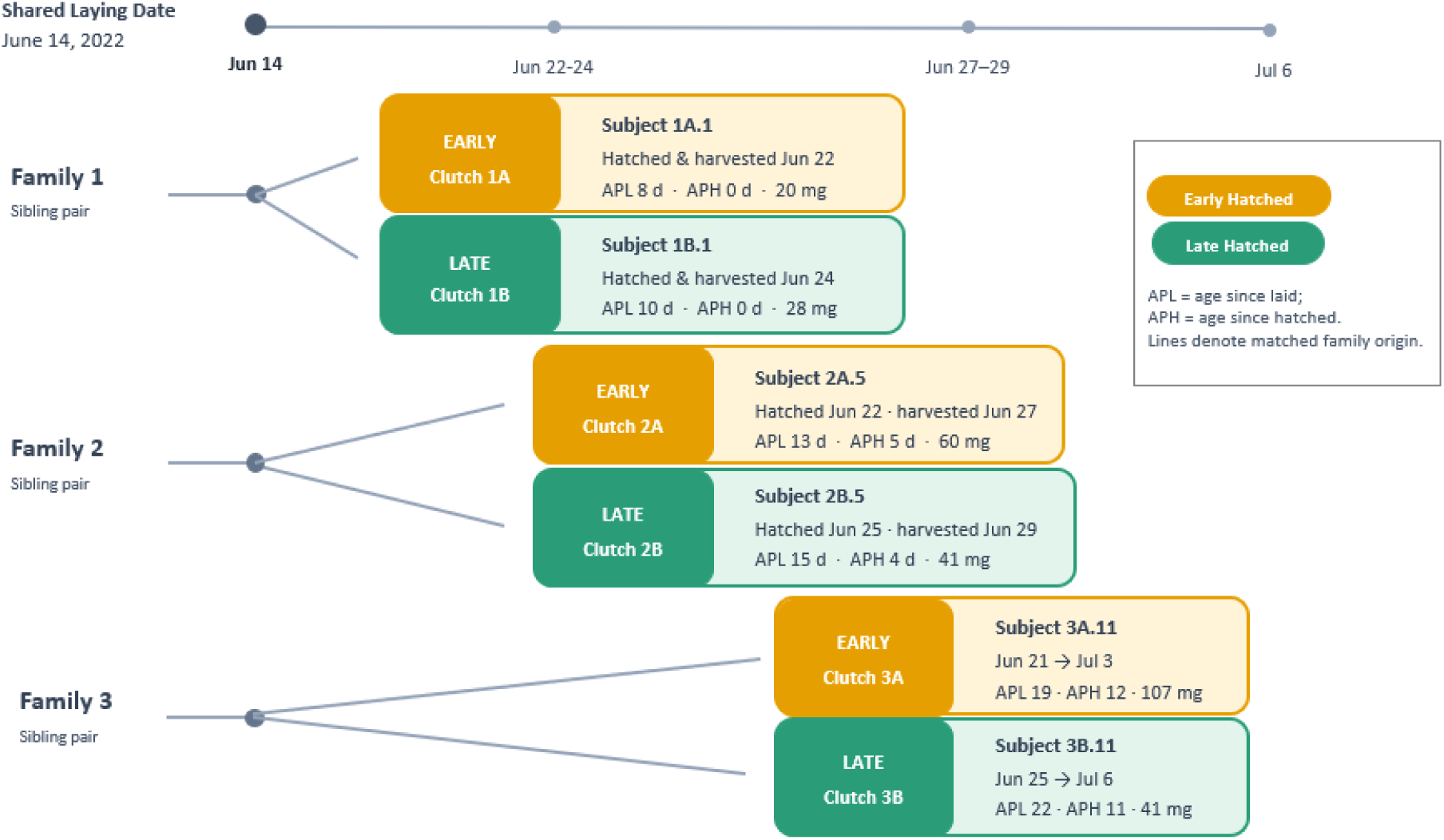
Origins, Sampling Timeline, and Body Mass of RNA-seq Subjects. RNA-seq subjects were sourced from three matched family pairs. Two siblings of the same laying date were early-hatched (EH: N=3) and late-hatched (LH: N=3) for comparisons within shared family origin. Body mass did not significantly differ between EH and LH (paired *t*(2) = 1.19, *p* = .357; exact Wilcoxon signed-rank test, *p* = .50).

**Figure 3.**
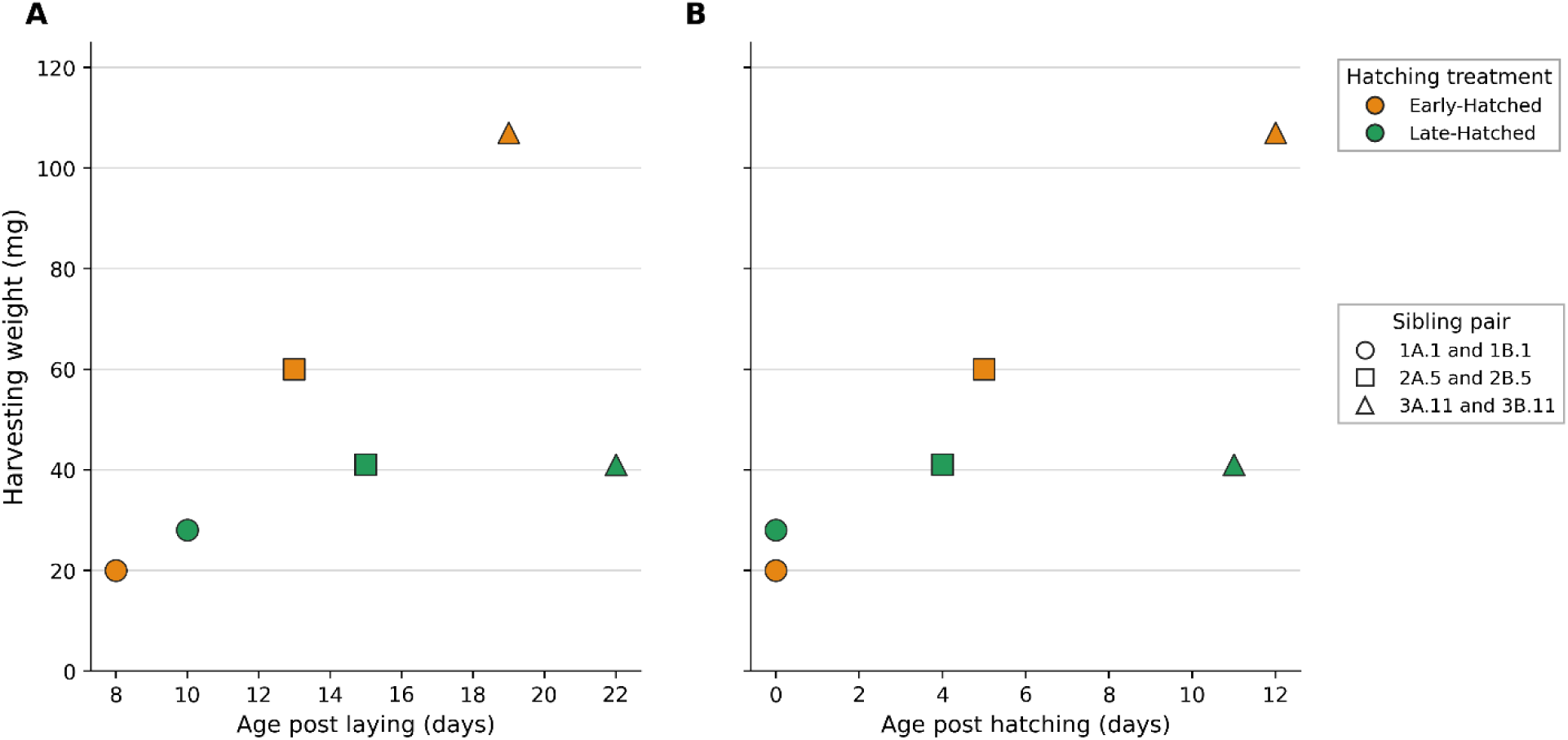
Individual RNAseq Subject Harvesting Weights by Chronological and Post-Hatch Age. Harvesting weight is shown for Early-Hatched (orange; *n* = 3) and Late-Hatched (green; *n* = 3) tadpoles. (**A**) Weight by age post laying (APL) and (**B**) Weight by age post hatching (APH) show identical patterns. Matching symbols identify sibling pairs: circles, 1A.1 and 1B.1; squares, 2A.5 and 2B.5; triangles, 3A.11 and 3B.11. Individual values were: 1A.1 (Early-Hatched), 20 mg, APL = 8 days, APH = 0 days; 1B.1 (Late-Hatched), 28 mg, APL = 10 days, APH = 0 days; 2A.5 (Early-Hatched), 60 mg, APL = 13 days, APH = 5 days; 2B.5 (Late-Hatched), 41 mg, APL = 15 days, APH = 4 days; 3A.11 (Early-Hatched), 107 mg, APL = 19 days, APH = 12 days; and 3B.11 (Late-Hatched), 41 mg, APL = 22 days, APH = 11 days.

***v. Body Size and Shape Analysis –*** Morphometric analyses were conducted in 67 tadpoles (EH = 32, LH = 35) with complete linear measurements; all were preserved in MEMFA and stored in 70% EtOH. Each tadpole was placed on a slide, photographed against a background of 1mm graph paper, and measured using ImageJ software (Schneider et al., 2012), by trained observers who were blind to treatment conditions. Measures included total tadpole length (TTL), body length (BL), body height (BH), tail-muscle height (TMH), and tail-fin height (TFH), in keeping with methods described by Touchon and Wojdak (2014) [Figure 4]. Harvesting mass was available for 61 of these animals (27 EH, 34 LH).

**Figure 4:**
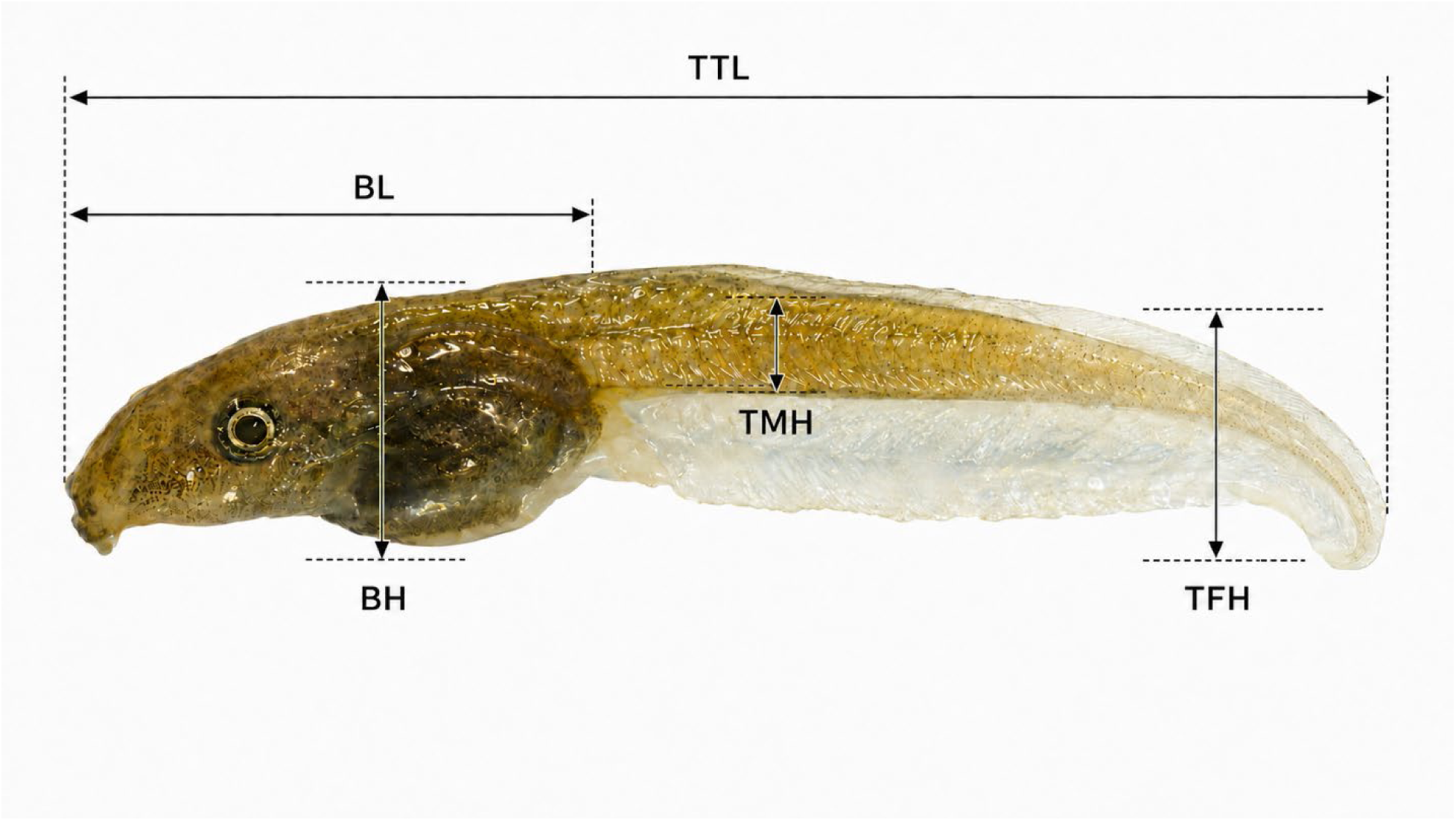
Red-Eyed Treefrog Tadpole Morphological Measurements. Lateral morphometric measurements of an *Agalychnis callidryas* tadpole from our study. Subject 7A.6: total tadpole length (TTL) = 26.813 mm; body length (BL) = 9.623 mm; body height (BH) = 3.875 mm; tail-muscle height (TMH) = 2.273 mm; and tail-fin height (TFH) = 4.506 mm.

### IV. Transcriptomic Studies

#### i. RNA Extraction and sequencing

Total RNA was extracted from whole tadpoles, per manufacturer’s instructions, using the Qiagen RNeasy Plus Universal mini kit (Qiagen, Hilden, Germany). RNA sample extracts were quantified using Qubit 2.0 Fluorometer (Life Technologies, Carlsbad, CA, USA). Then, RNA integrity was evaluated using Agilent TapeStation 4200 (Agilent Technologies, Palo Alto, CA, USA).

RNA sequencing libraries were prepared, per manufacturer’s instructions, using the NEBNext Ultra II RNA Library Prep for Illumina (NEB, Ipswich, MA, USA). Briefly, mRNAs were initially enriched with Oligod(T) beads. Enriched mRNAs were fragmented for 15 minutes at 94°C. First strand and second strand cDNA were subsequently synthesized. cDNA fragments were end repaired and adenylated at 3’ends, and universal adapters were ligated to cDNA fragments, followed by index addition and library enrichment by PCR with limited cycles. The sequencing libraries were validated on the Agilent TapeStation (Agilent Technologies, Palo Alto, CA, USA), and quantified by using Qubit 2.0 Fluorometer (Invitrogen, Carlsbad, CA) as well as by quantitative PCR (KAPA Biosystems, Wilmington, MA, USA).

The sequencing libraries were clustered on an Illumina NovaSeq flowcell. After clustering, the flowcell was loaded on the Illumina NovaSeq 6000 instrument according to manufacturer’s instructions. The samples were sequenced using a 2x150bp Paired End (PE) configuration. Image analysis and base calling were conducted by the Control software. Raw sequence data (.bcl files) generated by the sequencer were converted into fastq files and de-multiplexed using Illumina’s bcl2fastq 2.17 software. One mismatch was allowed for index sequence identification. Read outputs ranged from 104.0 – 133.6 million read pairs per sample. Raw reads were trimmed for quality and adapter contamination using Trimmonatic v. 0.36 (Bolger et al., 2014) and skewer v. 0.2.2 (Jiang et al., 2014) under default parameters. Trimmed read quality was evaluated using FastQC (Andrews, 2010).

#### ii. Transcriptome Assembly and Annotation

Because a reference genome is not available for *A. callydrias*, we assembled and annotated a de novo transcriptome from the RNAseq reads. We pooled reads from all individuals and generated a transcriptome assembly using Trinity v. 2.15.1 (Grabherr et al., 2011). The raw assembly contained 2.05M transcripts corresponding to 1.3M “genes”. We collapsed redundant sequences (>98% identical) using CD-HIT v. 4.7 (Fu et al., 2012), which resulted in 1.9M transcripts from 1.3M genes. Considering transcriptome assemblies produced from high-depth sequencing efforts are expected to contain a large number of sequences derived from pre-mRNAs, non-coding RNAs, assembly artifacts, and exogenous (i.e. contaminant) transcripts (Raghavan et al., 2022), we queried our transcripts with blastx v. 2.12.0+ (Camacho et al., 2009) against the SwissProt database (release 2024_01) and retained only transcripts whose best blast hit was a chordate protein. The filtered assembly contained 208,656 transcripts from 106,847 genes. To evaluate the degree to which this reduced set of sequences represented the mRNAs transcribed by *A. callydrias*, we ran BUSCO v.5.5 (tetrapoda_odb10 set; Manni et al., 2021; Simão et al., 2015) and mapped all reads against the filtered transcriptome using bwa v. 0.7.17-r1188 (Li, 2013). Ninety-eight percent of the BUSCO genes were found (97.0% complete, 1.1% fragmented) and 87.6% of the reads mapped to the filtered assembly, indicating adequate representation of the *A. callydrias* transcriptome. We therefore restricted downstream analyses to this reduced set of sequences.

To annotate the transcriptome, we used TransDecoder v. 5.7.1 (https://github.com/TransDecoder/TransDecoder) to identify likely coding regions with a minimum length of 50 AA, and Trinotate v. 4.0.2 (Bryant et al., 2017) to compile and integrate the results from multiple homology and orthology searches, and functional predictions. The SwissProt database was queried with blastp and blastx, the Rfam database (release 14.10) was searched with Infernal (cmscan) v. 1.1.5 (Nawrocki and Eddy, 2013), and the Pfam database v. 37 was searched with HMMER (hmmsearch) v. 3.1b2 (http://hmmer.org/). Signal peptides were predicted with signalp6 (Teufel et al., 2022), and transmembrane helices were predicted using tmhmm v.2.0 (Krogh et al., 2001). Finally, orthology predictions were done using eggNOG-mapper v. 2.1.12 (Cantalapiedra et al., 2021), based on version 5 of the eggNOG database (Huerta-Cepas et al., 2019), and using Diamond (Buchfink et al., 2021) for sequence searches.

### V. Statistical Analyses

#### i. Differential Gene Expression Analysis

Differential expression analysis was performed between the EH (N = 3) and LH (N = 3) groups using negative binomial generalized linear models. We used the align_and_estimate_abundance.pl script distributed with Trinity to map the trimmed reads from each sample to the unfiltered transcriptome assembly using bowtie2 v. 2.4.4 (Langmead and Salzberg, 2012), estimate transcript abundances using RSEM v. 1.3.3 (Li and Dewey, 2011), and average read counts for each Trinity “gene”. We used the unfiltered transcriptome assembly for read mapping to avoid reads from removed transcripts being incorrectly aligned to retained transcripts. We generated a read count matrix for all samples using the abundance_estimates_to_matrix.pl script, also distributed with Trinity, and removed counts for all genes that did not match chordate sequences in SwissProt, as well as those with less than 10 reads for at least three samples. To get a general idea of variation across our samples, we variance-stabilized the filtered counts matrix using the vst() function in the R package DESeq2 (version 1.38.3; Love et al., 2014), and performed PCA with the prcomp() function in R. Finally, we used DESeq2 to fit negative binomial models on each gene’s read counts, with treatment (EH vs. LH) as the focal predictor, clutch as a covariate. Significance for the effect of treatment on read counts was assessed with Wald’s test, and p-values were adjusted using the false discovery rate (FDR) method (Benjamini and Hochberg, 1995). Genes with adjusted p<0.05 and at least a two-fold shift in expression (i.e. log_2_(fold-change) > 1 or < -1) were considered differentially expressed between treatments.

#### ii Gene Ontology Analysis

Gene ontology (GO) terms for each gene were extracted from the Trinotate annotation using the extract_GO_assignments_from_Trinotate_xls.pl script distributed with Trinotate. We used the R package topGO v. 2.58.0 (Alexa and Rahnenführer, 2026) to identify statistically overrepresented GO terms from our differentially expressed gene set relative to the set of GO terms associated with all tadpole expressed genes using Fisher’s exact test. We collapsed redundant GO terms from overrepresented sets by calculating semantic similarity scores using the godata() and mgoSim() func 1tions from the R package GOSemSim v. 1.30.2, applying hierarchical clustering using the hclust() function from the R package stats v. 3.6.2, and selecting the GO term with the lowest weighted Fisher score from each cluster.

#### iii. Body Mass

Among 179 unique EH/LH tadpoles (80 EH, 99 LH), six lacked harvesting mass, leaving 173 observations for body-mass analyses (75 EH, 98 LH). Age post-laying (APL) was calculated as the interval between laying and harvesting, and age post-hatching (APH) as the interval between hatching and harvesting. Treatment differences in APL and APH sampling distributions were evaluated using two-sided variance-ratio tests, median-centered Levene tests, and Welch two-sample *t* tests.

Each tadpole contributed one harvesting mass, so age slopes describe cross-sectional mass–age associations. Separate ordinary least-squares ANCOVA models for APL and APH included age, treatment, and their interaction, with EH as the reference treatment. The age-by-treatment coefficient therefore estimated the LH−EH difference in slope.

Conventional inference was supplemented with HC3 heteroscedasticity-consistent covariance estimates; contrast tests and 95% confidence intervals used the residual degrees of freedom and the t distribution. The full APH model used all 173 mass-valid observations, and a common-range analysis restricted to 0–17 days APH used 164 observations (66 EH, 98 LH). These primary linear mass models did not include family effects.

Age-dependent changes in the APH association were also evaluated using a continuous piecewise-linear model with fixed knots at 7 and 14 days. The model included treatment, APH, the hinge terms (APH−7)h and (APH−14)h, and all treatment-by-age-term interactions, where (x)h = max(x, 0). Linear contrasts estimated treatment-specific slopes and LH−EH slope differences over 0–7, >7–14, and >14 days APH. These intervals corresponded to observed integer-day windows of 0–7, 8–14, and ≥15 days.

Separate Welch comparisons of mean mass and age-by-treatment slope models within these windows were treated as secondary analyses. Conventional and HC3 sensitivity results were considered together, with discrepancies explicitly acknowledged. The joint weekly-segment likelihood comparison contrasted a model containing window, APH, treatment, and all interactions with a reduced model containing window, APH, and their interaction. A single residual variance was estimated across windows; the models differed by six mean-model coefficients. The reported likelihood-ratio p-value used a chi-square approximation with six degrees of freedom. Additional secondary comparisons used the RNAseq-centered windows of 0–12 and ≥13 days APH. Window-specific and targeted follow-up p-values were unadjusted for multiplicity.

Three control specimens decapsulated on crushed ice (without hatching) were excluded from EH–LH model fitting and statistical comparisons. Primary GAM predictions were not anchored to control mass. For Figures 6a and 7 only, illustrative trajectories were constructed by propagating the estimated raw-mass linear slopes from the control mean of 17.33 mg. This display convention did not enter the regression fits or alter the estimated linear slopes.

#### iv. Body Size and Shape

TTL, BL, BH, TMH, and TFH were natural-log transformed, and each animal’s mean log dimension defined its log geometric-mean size. APH was centered at the morphometry-sample mean of 11.1343 days, and the square of this centered age was calculated. The size model included treatment, centered APH, centered APH^2^, both treatment-by-age terms, and Family of Origin as a fixed blocking factor. Partial F tests evaluated the treatment main effect and both interactions jointly (3 df) and the two interactions jointly (2 df). Predictions at APH 10 were averaged equally over the nine family levels on the log-size scale and then exponentiated; confidence limits used the full-model residual degrees of freedom.

For shape, each log dimension was centered by subtracting that animal’s log geometric-mean size. Principal Component Analysis (PCA) of the five centered log dimensions yielded four nonredundant shape PCs, as expected because size centering imposes a sum-to-zero constraint and reduces the five-dimensional shape space to rank four. The four PCs were analyzed jointly with treatment, mean-centered APH, treatment × APH, log geometric-mean size, and Family of Origin. Pseudo-F statistics were based on the summed residual sums of squares across PCs. A two-degree-of-freedom test evaluated the joint treatment profile (treatment and treatment × APH), and a one-degree-of-freedom test evaluated treatment × APH. Significance was estimated using 9,999 Freedman–Lane residual permutations restricted within Family of Origin. Separate reduced models were used for the joint treatment profile and interaction hypotheses; the same residual-row permutation was applied to all four PC responses. Seeds were 20261029 and 20261030, respectively. The permutation p-value was calculated as one plus the number of permuted statistics at least as large as the observed statistic, divided by 10,000. Secondary outcomes were log(BL/TTL), log(BH/TTL), log(TMH/TTL), log(TFH/TTL), and log(TFH/TMH), modeled with the same treatment, APH, treatment × APH, log-size, and family covariates.

#### v. Comparison with Published *Agalychnis callidryas* Size and Mass Data

To place body-size measurements from our laboratory-reared tadpoles in the context of prior work on *A. callidryas*, we compared total tadpole length (TTL) and wet mass descriptively with published values spanning hatching through later larval development. Because these studies differed in hatching age, rearing environment, density, experimental treatment, developmental age at measurement, and methods of estimating or measuring mass, the published values were used as comparative benchmarks rather than as normative criteria for inclusion or exclusion of observations. Where the original publication reported numerical values, those values were used directly; where only graphical data were available, approximate values were taken from published figures.

Published measurements indicate relatively little variation in body size around hatching. For Late-Hatched (LH) tadpoles, Asquith and Vonesh (2012) reported approximately 11.13 mm total length and 10.08 mg wet mass at hatching or six days post-oviposition. In two experiments, McCoy et al. (2015) reported six-day LH hatchlings averaging approximately 11.75 mm and 11.78 mg, and 11.54 mm and 11.18 mg, respectively. Warkentin (1999) reported an estimated total length of approximately 12.5 mm and wet mass of 14.06 mg for LH tadpoles hatching at seven days. Early-Hatched (EH) tadpoles were correspondingly smaller: values estimated from Warkentin (1999) were approximately 10.5 mm and 8.53 mg for five-day EH hatchlings, while Touchon et al. (2015) reported approximately 9.5 mm and 6.40 mg for four-day EH hatchlings. Wet-mass measurements reported by Rogge and Warkentin (2008) provide a similar scale, with five-day embryos weighing 13.7 ± 0.06 mg at hatching and 15.2 ± 0.4 mg after approximately 1.8 h in water.

Against this published range, linear measurements from our study were broadly comparable at hatching, particularly for LH tadpoles. LH animals measured at hatching had TTL values of approximately 12–13 mm, overlapping the approximately 11–13 mm values reported by Asquith and Vonesh (2012), McCoy et al. (2015), and Warkentin (1999). In contrast, their recorded wet masses, approximately 28–37 mg, were substantially higher than the approximately 10–15 mg hatchling masses reported in these studies. EH hatchling masses in our dataset were also comparatively high relative to the approximately 6–9 mg values estimated or reported for four- to five-day EH hatchlings by Warkentin (1999) and Touchon et al. (2015).

Differences between our measurements and published values became less uniform with increasing post-hatching age. For example, Asquith and Vonesh (2012) estimated an LH mass of approximately 46 mg at nine days post-hatching. McCoy et al. (2015) reported approximately 20.7 mm and 59.7 mg, and 24.2 mm and 94.0 mg, for LH tadpoles seven days post-hatching in two different experimental contexts. Warkentin (1999) reported approximately 20 mm and 54.2 mg for EH and LH tadpoles at 14 days post-oviposition, corresponding to approximately nine and seven days post-hatching, respectively. Touchon et al. (2015) reported approximately 26.5 mm and 121.5 mg for EH tadpoles eight days post-hatching and approximately 25 mm and 102.8 mg for LH tadpoles six days post-hatching. Thus, by approximately one week after hatching, wet masses in our study overlapped more substantially with the broad range represented in previous studies, although our tadpoles were often shorter than animals of comparable post-hatching age in some published datasets.

Published later-larval measurements further illustrate the strong context dependence of size-at-age. Touchon et al. (2013) reported tadpoles of approximately 42 mm and 456 mg at 15 days post-hatching, with little remaining difference between EH and LH groups. Touchon et al. (2015) similarly reported EH tadpoles of approximately 43 mm and 488 mg at 18 days post-hatching. Under different competition conditions, Bouchard et al. (2016) reported that LH tadpoles reached approximately 45 mm and 700 mg after about 12 days under low competition but required approximately 20 days to reach the same size under high competition. These differences among published studies demonstrate that larval size-at-age is sensitive to rearing and ecological conditions and therefore provide context for, rather than a fixed standard against which to judge, the developmental trajectories observed in our colony.

Body mass and linear morphometrics were therefore treated as complementary rather than interchangeable measures of phenotype. Wet mass of small aquatic larvae can be affected by hydration and residual surface water at weighing, whereas total length may be influenced by specimen position, curvature, or tail condition. In addition, variation among studies in rearing density, food availability, temperature, hatching age, and other environmental conditions can alter size-at-age. Accordingly, the mass–age analyses in the present study were used to characterize cross-sectional differences between EH and LH tadpoles maintained and sampled under the same experimental conditions, rather than to establish a species-typical growth trajectory. TTL provides a directly interpretable measure of absolute linear size that can be compared with prior *A. callidryas* studies, while the multivariate morphometric analyses integrate TTL with body and tail dimensions to characterize overall size and proportional body shape.

#### vi. Post-Hatching Growth-Curves

Mass–age trajectories were modeled using Gaussian generalized additive models of natural-log harvesting mass. The full APH model included treatment, treatment-specific thin-plate regression spline smooths of APH (basis dimension k = 4 for each treatment), and a random-intercept smooth for Family of Origin. Smoothing parameters were estimated by maximum likelihood. The reduced model retained the treatment main effect and family random effect but used a common APH smooth. Both models used the same 173 mass-valid EH/LH animals (75 EH, 98 LH). Overall trajectory difference was evaluated using an approximate likelihood-ratio statistic, calculated as twice the difference in model log likelihoods; its p-value used a chi-square approximation with degrees of freedom equal to the difference in effective model degrees of freedom.

An equivalent treatment-specific GAM using APL generated the chronological-age curves. APH curve and derivative contrasts were restricted to shared support of 0–17 days; 164 animals (66 EH, 98 LH) were observed within that domain, although all 173 informed the model fit. APL curve comparisons were restricted to the shared 10–25-day domain. To avoid comparing treatments outside common data support, between-treatment curve and derivative contrasts were evaluated only over 0–17 days APH, which is the age range observed in both treatments. Predictions set the family random-effect contribution to zero. Exponentiating the fitted log-mass prediction produced an unanchored conditional-median mass curve. First and second derivatives of this curve, in mg/day and mg/day², described changes in cross-sectional fitted mass with age.

Derivatives were calculated from central finite differences of the GAM linear-predictor matrix with a step of 0.001 day and the exponential chain rule.

Approximate simultaneous 95% bands were constructed separately for the EH−LH first- and second-derivative contrasts across a 0.01-day grid from APH 0 to 17. Delta-method gradients and the smoothing-uncertainty-corrected coefficient covariance were used with 20,000 Gaussian coefficient-error simulations (seed 20261031). For each derivative order, the 95th percentile of the maximum absolute standardized contrast error over the age grid supplied the simultaneous critical value. A band excluding zero identified a supported difference under that model; a band including zero did not establish equivalent rates. Sensitivity analyses used k = 6 and k = 8 and repeated the k = 4 APH GAM after excluding the specified five-record QC batch.

#### vii. Statistical Packages

Standard analyses were conducted in SPSS Statistics 32 Software (IBM, Inc.). Mass, size, shape, and GAM analyses were conducted in R version 4.3.3 using base and recommended packages (R Core Team, 2024; packages car, lmtest, sandwich, emmeans, RRPP, and mgcv version 1.9-1). HC3 covariances, linear contrasts, equal-family size predictions, and Freedman–Lane permutation tests were implemented in the accompanying analysis script. Exact package versions and session information accompany the reproducibility files. Regression slopes describe cross-sectional mass– age associations; intercepts are model predictions at age zero, and residual SD quantifies scatter around the fitted regression.. Figures were produced in GraphPad Prism 11.1.0 (GraphPad Software, Inc.) or Microsoft Office. Reported findings include the slope (rate of change), intercept (starting value, where the line crosses the y-axis), and residual SD (Sy.x: the typical prediction error of the regression line). Transcriptome analyses used the packages reported in sections IV and V.

## RESULTS

### Growth Trajectory: Age Post Laying

Body mass was positively associated with age post-laying (APL) in both groups. EH tadpoles (N = 75) had an estimated cross-sectional slope of 3.36 mg/day (SE = 0.43, 95% CI [2.51, 4.21]), F(1, 73) = 61.99, p < .001, R² = .459, residual SD = 17.47 mg. LH tadpoles (N = 98) had an estimated slope of 6.24 mg/day (SE = 1.43, 95% CI [3.40, 9.08]), F(1, 96) = 18.97, p < .001, R² = .165, residual SD = 55.22 mg. The LH−EH slope difference was 2.88 mg/day (95% CI [−0.16, 5.91]), t(169) = 1.87, p = .063. HC3 inference was slightly more favorable (p = .046), so evidence for an APL slope difference was sensitive to the variance estimator rather than robustly null. The full interaction model explained 21.2% of mass variance (R² = .212). [Figure 5a].

**Figure 5a.**
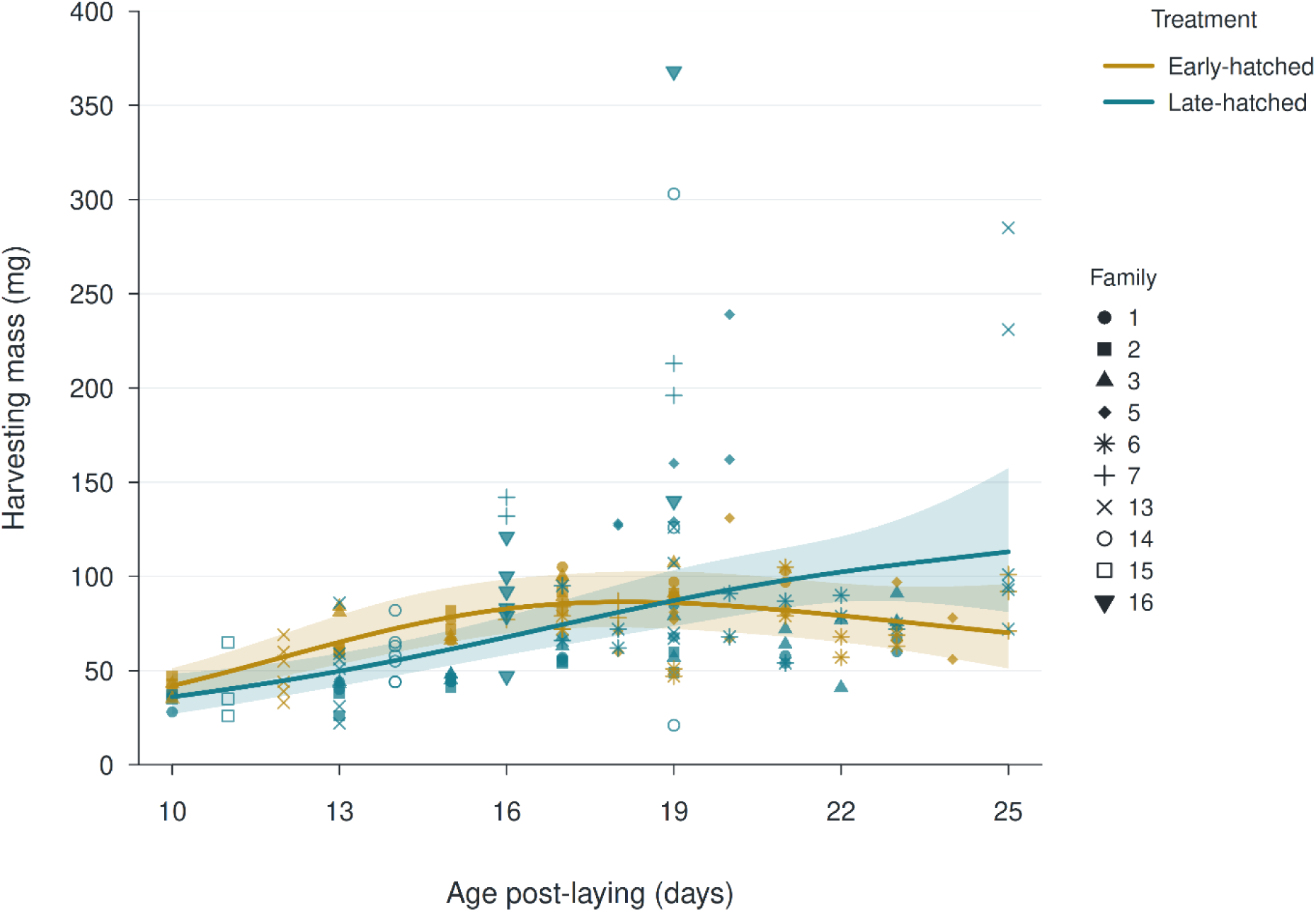
Age Post-Laying and Mass-Growth Trajectories of Early- and Late-Hatched Tadpoles. Harvesting mass and fitted mass–APL relationships in early- and late-hatched tadpoles. Raw harvesting mass is plotted against age post-laying (APL). Curves are unanchored conditional-median predictions from a Gaussian GAM of log mass with treatment-specific thin-plate smooths (k = 4), maximum-likelihood smoothing selection, and a family random-intercept smooth; the family contribution is set to zero for prediction. Ribbons show pointwise 95% intervals. Curve comparisons are displayed over the shared 10–25-day APL domain. The fitted EH curve peaks near 18.13 days APL, and the treatment curves cross once within this domain, near 18.83 days APL; EH predicted mass is greater before this crossing and LH predicted mass is greater afterward. Separate descriptive linear regressions over each treatment’s full observed APL range give slopes of 3.36 ± 0.43 SE mg/day for EH (n = 75; 95% CI [2.51, 4.21]; F(1, 73) = 61.99, p < .001, R² = .459, residual SD = 17.47 mg) and 6.24 ± 1.43 SE mg/day for LH (n = 98; 95% CI [3.40, 9.08]; F(1, 96) = 18.97, p < .001, R² = .165, residual SD = 55.22 mg).

### Growth Trajectory: Age Post Hatching

When aligned by age post-hatching (APH), body mass was positively associated with age in both groups, but the cross-sectional association was steeper in LH tadpoles.

Across the full observed APH range, the EH mass–APH slope (N = 75) was 2.56 ± 0.39 SE mg/day (95% CI [1.79, 3.34]), F(1, 73) = 43.78, p < .001, R² = .375, residual SD = 18.78 mg; the fitted EH equation was Mass = 44.24 + 2.56×APH (intercept 95% CI [35.35, 53.12]). The LH mass–APH slope (n = 98) was 7.05 ± 1.09 SE mg/day (95% CI [4.88, 9.22]), F(1, 96) = 41.54, p < .001, R² = .302, residual SD = 50.49 mg; the fitted LH equation was Mass = 19.78 + 7.05×APH (intercept 95% CI [−2.36, 41.93]). The LH−EH slope difference was 4.49 mg/day (95% CI [2.12, 6.85]), t(169) = 3.75, p = .00024; HC3 p = .00037. The full interaction model explained 32.3% of mass variance (R² = .323).

When analysis was restricted to the shared 0–17-day APH range, the conclusion was unchanged (n = 164: 66 EH, 98 LH): LH−EH = 3.63 mg/day (95% CI [0.94, 6.32]), t(160) = 2.67, p = .0084; HC3 p = .0049, R² = .333. [Figure 5b].

**Figure 5b:**
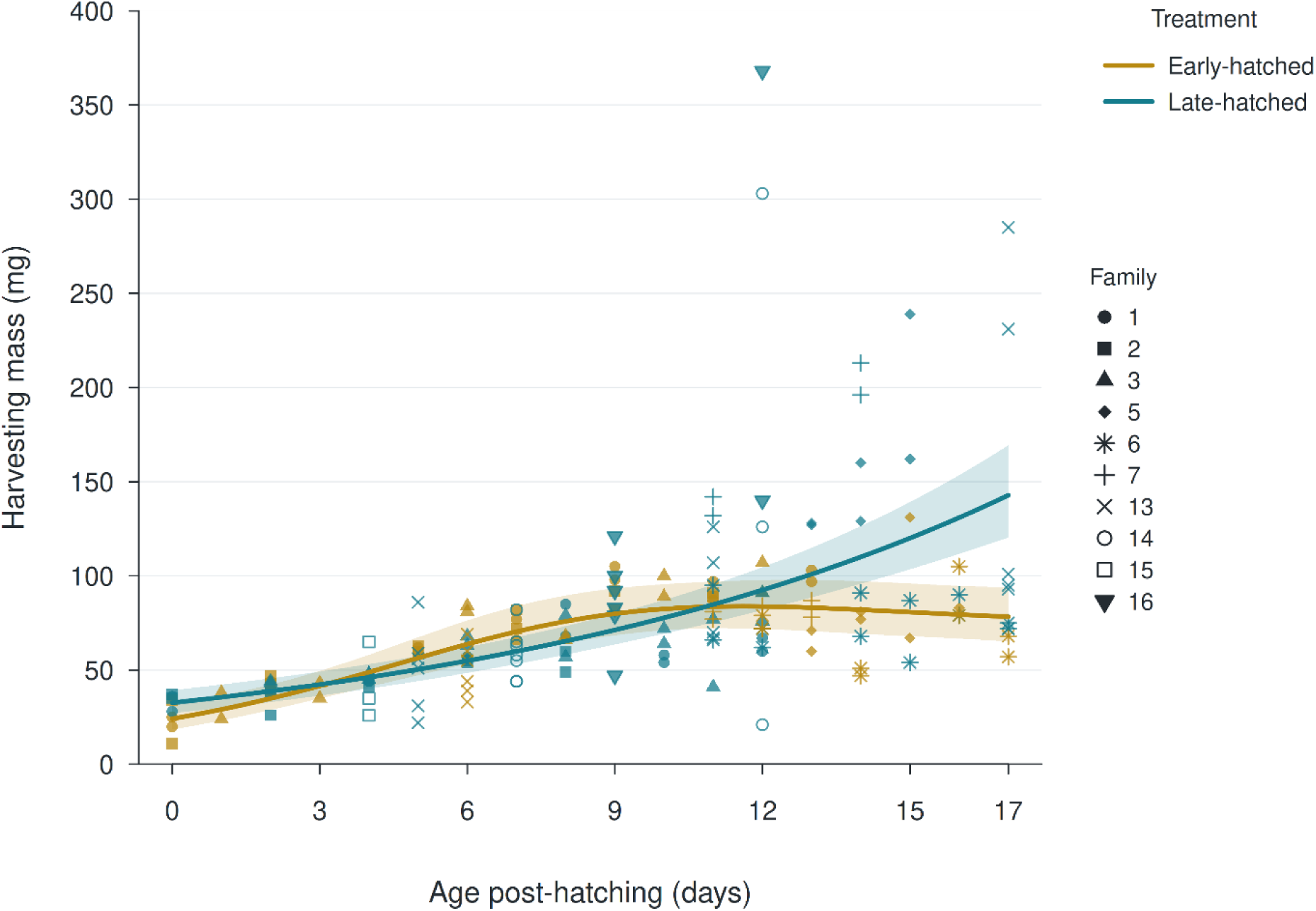
Age Post Hatching and Mass-Growth Trajectories of Early- and Late-Hatched Tadpoles. Harvesting mass and fitted mass–APH relationships in early- and late-hatched tadpoles. Raw harvesting mass is plotted against age post-hatching (APH). Curves are unanchored conditional-median predictions from the cleaned-data Gaussian GAM of log mass, with treatment-specific thin-plate smooths (k = 4), maximum-likelihood smoothing selection, and the family random-effect contribution set to zero. Ribbons show pointwise 95% intervals. The model uses all 173 mass-valid animals but displayed between-treatment comparisons are restricted to shared support of 0–17 APH. The EH curve peaks near 11.61 days APH; the treatment mass curves cross near 3.22 and 10.83 days APH, with EH predicted mass greater between the crossings and LH predicted mass greater outside them. Control masses do not anchor these predictions. Separate full-range linear regressions give EH slope = 2.56 ± 0.39 SE mg/day (n = 75; 95% CI [1.79, 3.34]; F(1, 73) = 43.78, p < .001, R² = .375, residual SD = 18.78 mg) and LH slope = 7.05 ± 1.09 SE mg/day (n = 98; 95% CI [4.88, 9.22]; F(1, 96) = 41.54, p < .001, R² = .302, residual SD = 50.49 mg).

### Growth by Weekly Segments

Raw harvesting mass and segment-specific mass–APH associations were examined across 0–7, 8–14, and ≥15 days post-hatching. During days 0–7, mean mass did not differ between EH tadpoles (n = 28, M = 50.07 mg, SD = 19.98) and LH tadpoles (n = 39, M = 48.26 mg, SD = 14.40), Welch p = .684. Within this window, the estimated slope was 6.45 mg/day for EH and 3.93 mg/day for LH. The conventional slope contrast was borderline but did not meet α = .05 (LH−EH = −2.52 mg/day, t(63) = −2.00, p = .0502), whereas the HC3 sensitivity test was significant (p = .026).

During days 8–14, EH tadpoles (n = 30, M = 82.63 mg, SD = 15.03) were lighter on average than LH tadpoles (n = 45, M = 101.98 mg, SD = 64.74), although the Welch comparison was not significant (p = .059). The EH slope was −2.48 mg/day and the LH slope was 13.44 mg/day; the slopes differed (LH−EH = 15.92 mg/day, t(71) = 2.56, p = .0126; HC3 p = .00044).

At ≥15 days, EH tadpoles (n = 17, M = 80.59 mg, SD = 19.47) were lighter than LH tadpoles (n = 14, M = 123.93 mg, SD = 74.23), with a borderline Welch test (p = .051). Neither group showed evidence of continued linear mass gain (EH = −2.30 mg/day; LH = −1.26 mg/day), and slopes did not differ (t(27) = 0.06, p = .955; HC3 p = .970). A joint likelihood-ratio comparison of treatment-specific segment trajectories was significant, χ²(6) = 23.14, p = .00075 [Figure 6a].

**Figure 6a:**
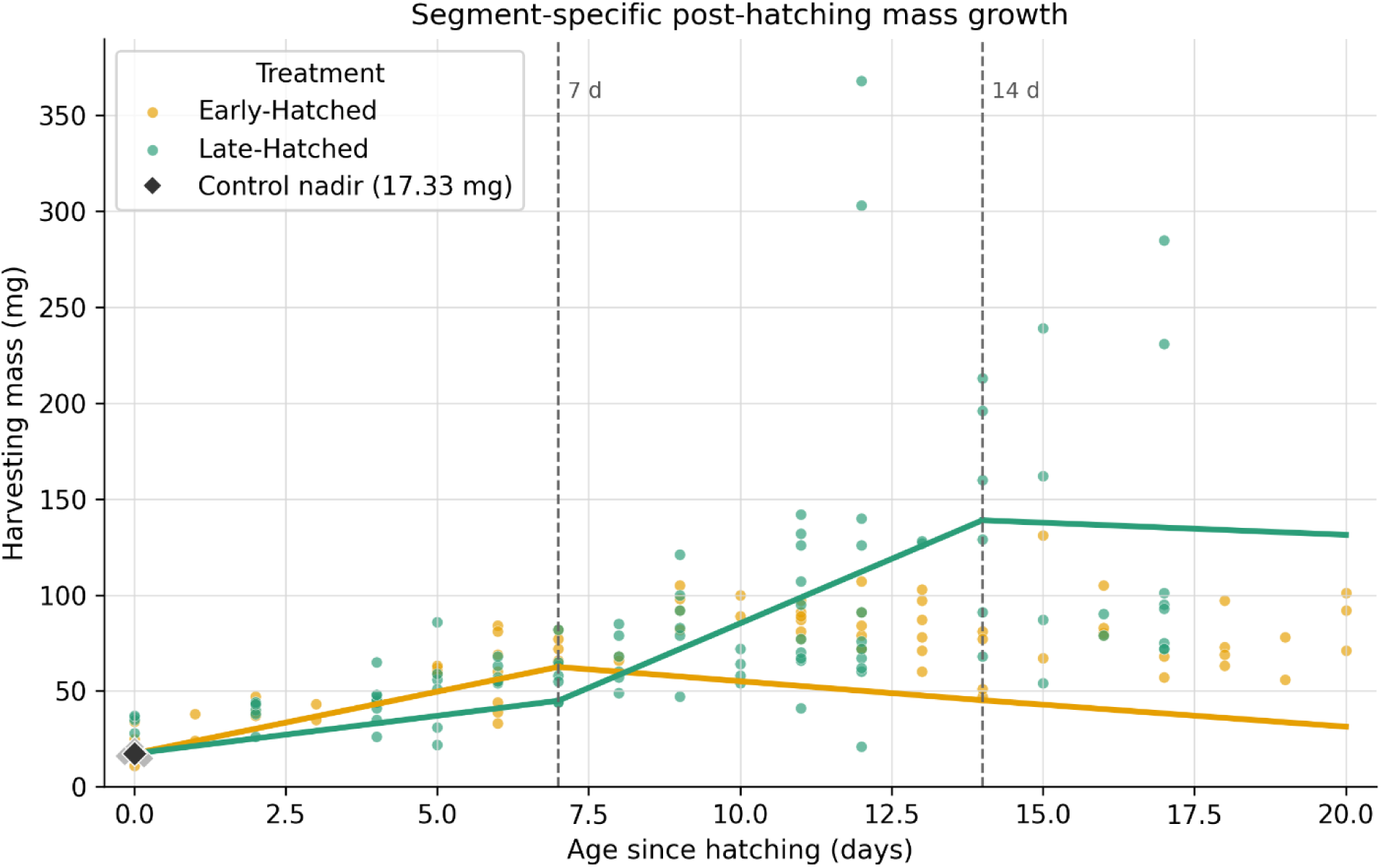
Time Segmented, Red-Eyed Treefrog Growth Patterns. Raw harvesting masses and illustrative trajectories are shown for EH (n = 75) and LH (n = 98) animals across the three APH periods. The displayed trajectories propagate the separately estimated raw-mass linear slopes from the pre-hatching control mean of 17.33 mg. This construction preserves the estimated segment slopes but does not represent fitted age-specific mean masses or observed individual growth paths. All regressions and tests exclude the controls and use the 173 mass-valid EH/LH animals. During days 0–7 (EH n = 28, LH n = 39), estimated slopes were 6.45 mg/day for EH and 3.93 mg/day for LH; the conventional difference narrowly missed α = .05, t(63) = −2.00 for LH−EH, p = .0502 (HC3 p = .026). During days 8–14 (EH n = 30, LH n = 45), EH = −2.48 mg/day and LH = 13.44 mg/day; slopes differed, t(71) = 2.56, p = .0126 (HC3 p = .00044). At ≥15 days (EH n = 17, LH n = 14), slopes were −2.30 and −1.26 mg/day and did not differ, p = .955. A joint likelihood-ratio comparison of treatment-specific segment trajectories was significant, χ²(6) = 23.14, p = .00075.

### Continuous Piecewise-Linear Model

A continuous piecewise-linear model with knots at 7 and 14 days APH (N = 173) was significant, F(7, 165) = 13.93, p < .001, and explained 37.1% of harvesting-mass variation (R² = .371). During days 0–7, the estimated EH slope was 7.66 mg/day, and the LH slope was 3.19 mg/day. The conventional LH−EH contrast was not significant (−4.47 mg/day, 95% CI [−11.87, 2.92], p = .234), but HC3 inference indicated a difference (95% CI [−6.98, −1.97], p = .00054); this first-week result was therefore sensitive to the covariance estimator. During days 8–14, the pattern reversed: EH = 1.18 mg/day and LH = 11.79 mg/day, with LH−EH = 10.62 mg/day (95% CI [4.59, 16.64], p = .00064; HC3 p = .00033). At ≥15 days, estimated slopes were negative in both groups (EH = −1.16 mg/day; LH = −5.61 mg/day), and the slope difference was not significant (LH−EH = −4.46 mg/day, 95% CI [−17.27, 8.36], p = .493; HC3 p = .697). [Figure 6b].

**Figure 6b:**
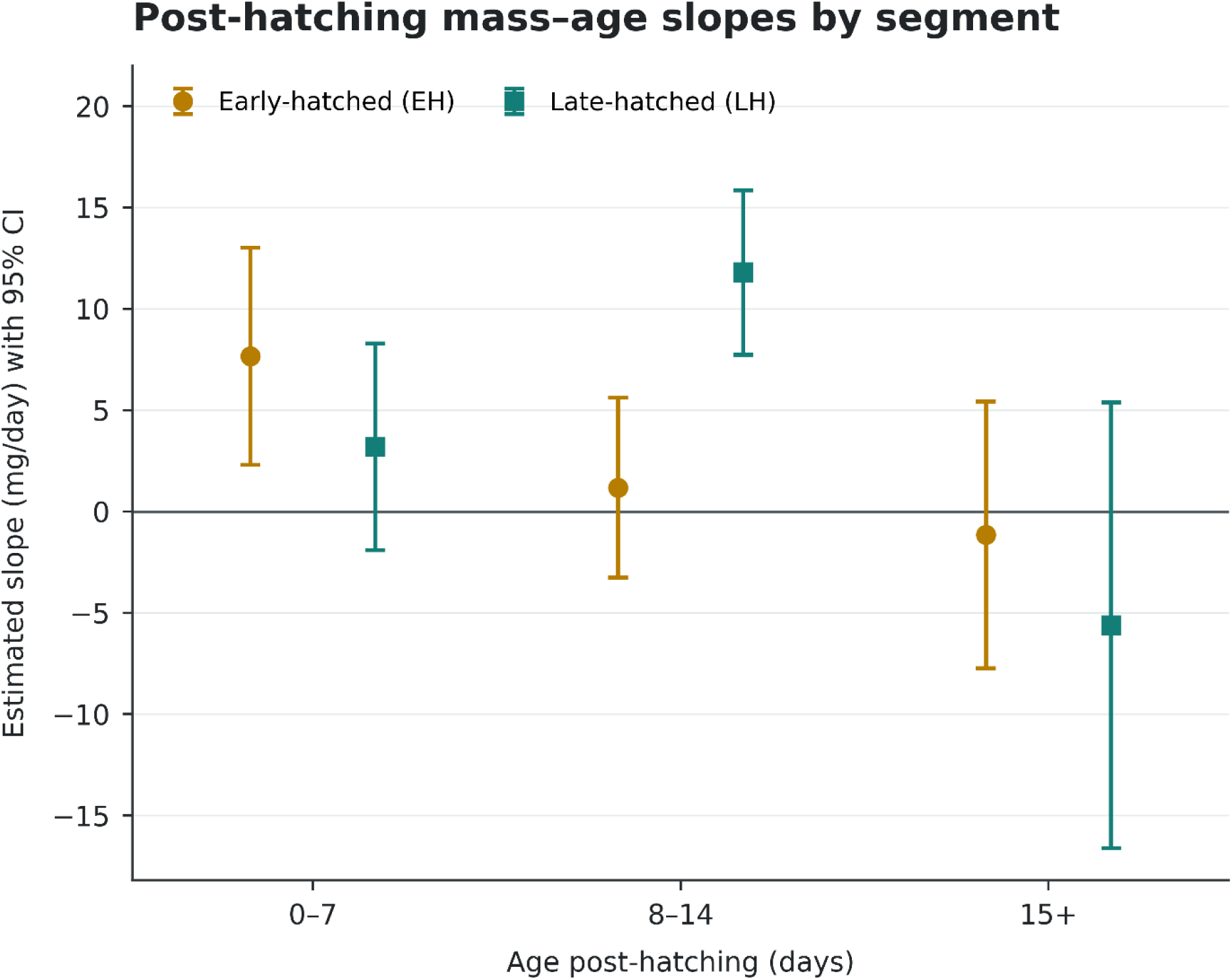
Slope of Post-Hatching Growth Data by Study Period. Points show treatment-specific slopes from the continuous piecewise-linear model with 95% confidence intervals (EH N = 75; LH N = 98). During days 0–7, the EH slope was +7.66 mg/day (95% CI [2.30, 13.03]) and the LH slope was +3.19 mg/day (95% CI [−1.90, 8.28]). The conventional LH−EH contrast was −4.47 mg/day, t(165) = −1.19, p = .234; HC3 95% CI [−6.98, −1.97], p = .00054. The HC3 test of the LH slope itself was p = .0012. During days 8–14, EH = 1.18 mg/day (95% CI [−3.27, 5.63]) and LH = 11.79 mg/day (95% CI [7.73, 15.85]); LH−EH = 10.62 mg/day, t(165) = 3.48, p = .00064. At ≥15 days, EH = −1.16 mg/day (95% CI [−7.74, 5.43]) and LH = −5.61 mg/day (95% CI [−16.61, 5.38]); LH−EH = −4.46 mg/day, t(165) = −0.69, p = .493.

### Follow-Up Data Quality Control and Sensitivity Analyses

The positive full-range APH slope difference and the joint weekly-segment result remained supported after excluding the five potentially anomalous LH specimens collected on June 21, 2023. In the full-range APH interaction model, the estimated LH−EH slope difference changed from 4.49 to 3.84 mg/day; conventional significance became more decisive (p = .00024 to p = .000040), while the HC3 test remained significant (p = .00037 in the full dataset; p = .0013 after exclusion). During days 8–14, the LH within-window slope decreased from 13.44 to 10.51 mg/day, but the conventional treatment contrast remained significant and became more precise, changing from t(71) = 2.56, p = .0126, to t(66) = 3.63, p = .00055 (HC3 p = .0020 after exclusion). The joint segment likelihood-ratio test changed from p = .00075 to p = .00016. Thus, the linear and pooled weekly-segment findings did not depend on this batch. Smaller conventional p-values after exclusion accompanied reduced residual dispersion rather than larger estimated treatment differences; the nonlinear timing estimates were evaluated separately in GAM sensitivity analyses.

### RNAseq Subject-Centered Growth Trajectories

Figure 7 highlights the six RNAseq animals within the full mass dataset and displays illustrative trajectories constructed from the separate 0–12 and ≥13 APH linear slopes, propagated from the 17.33-mg control reference. During the RNAseq-relevant 0–12 APH period, mass was positively associated with APH in both treatments, with estimated slopes of 5.53 mg/day for EH (95% CI [4.58, 6.49]) and 7.33 mg/day for LH (95% CI [4.44, 10.22]). The LH–EH slope difference was not significant (1.80 mg/day, 95% CI [−1.76, 5.36], p = .320; HC3 p = .358) and mean raw harvesting mass was also similar between groups (EH: 65.19 ± 24.75 mg, n = 48; LH: 70.51 ± 51.87 mg, n = 76; Welch p = .444). At ≥13 APH, LH tadpoles had significantly greater raw harvesting mass than EH tadpoles (129.41 ± 65.34 vs. 78.59 ± 18.92 mg; Welch p = .0017), but neither group exhibited a significant continuing mass–age slope (EH = −0.07 mg/day; LH = −4.11 mg/day), and the slopes did not differ (LH–EH = −4.04 mg/day, p = .612; HC3 p = .692).

### Growth Curves

*GAM Overall-Curve Test* — Post-hatching mass trajectories differed significantly between EH and LH tadpoles. A treatment-specific GAM provided substantially better fit than a common-shape model that allowed only an overall treatment difference (likelihood-ratio statistic = 23.21, Δdf = 1.93, p = 8.16 × 10⁻⁶), demonstrating an age-dependent difference in trajectory shape. The EH trajectory was distinctly nonlinear on the log-mass scale (edf = 2.72), whereas the LH trajectory was approximately linear on the log-mass scale (edf = 1.00) [Figure 8a]. *First-Derivative Results* - First-derivative contrasts with simultaneous 95% confidence bands showed that EH tadpoles grew significantly faster from 00–3.86 days APH, whereas LH tadpoles grew significantly faster from 9.89–17 days APH. The band included zero over 3.87–9.88 days APH, providing no detected difference during that interval under this specification. The fitted treatment growth rates crossed at 7.53 days APH, and EH slope reached zero at 11.61 days APH, marking the fitted EH mass maximum [Figure 8b].

**Figure 7:**
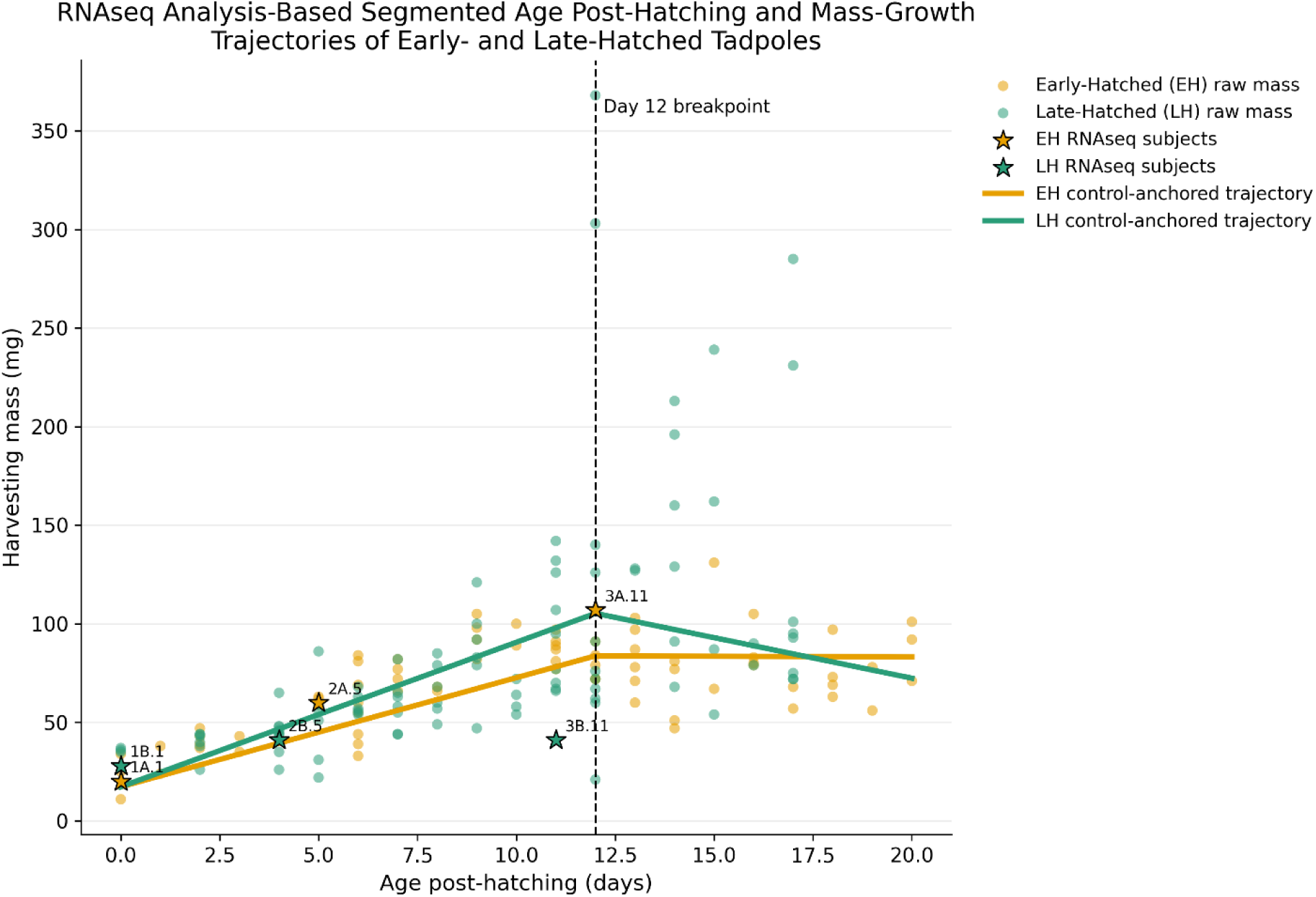
RNAseq Analysis Based Segmented Age Post-Hatching and Mass-Growth Trajectories of Early- and Late-Hatched Tadpoles. Figure shows all analyzable individual observations, highlights the six RNAseq animals, marks the Day 12 breakpoint, and overlays the EH and LH trajectories anchored to the 17.33-mg control mean. During the RNAseq-relevant 0–12 APH period, both EH and LH tadpoles increased significantly in mass, with estimated slopes of 5.53 mg/day for EH (95% CI [4.58, 6.49]) and 7.33 mg/day for LH (95% CI [4.44, 10.22]). The LH–EH slope difference was not significant (1.80 mg/day, 95% CI [−1.76, 5.36], p = .320; HC3 p = .358) and mean raw harvesting mass was also similar between groups (EH: 65.19 ± 24.75 mg, n = 48; LH: 70.51 ± 51.87 mg, n = 76; Welch p = .444). At ≥13 APH, LH tadpoles had greater raw harvesting mass than EH tadpoles (129.41 ± 65.34 vs. 78.59 ± 18.92 mg; Welch p = .0017), but neither group exhibited a significant continuing mass–age slope (EH = −0.07 mg/day; LH = −4.11 mg/day), and the slopes did not differ (LH–EH = −4.04 mg/day, p = .612; HC3 p = .692). Thus, the 0–12 APH window containing all six RNAseq animals had positive average linear mass–age slopes in both treatments without a detected between-treatment slope difference. This window-level result does not exclude age-dependent nonlinear differences within the interval; at ≥13 APH, the raw-mass contrast favored LH while ongoing linear slopes remained imprecisely estimated.

**Figure 8.**
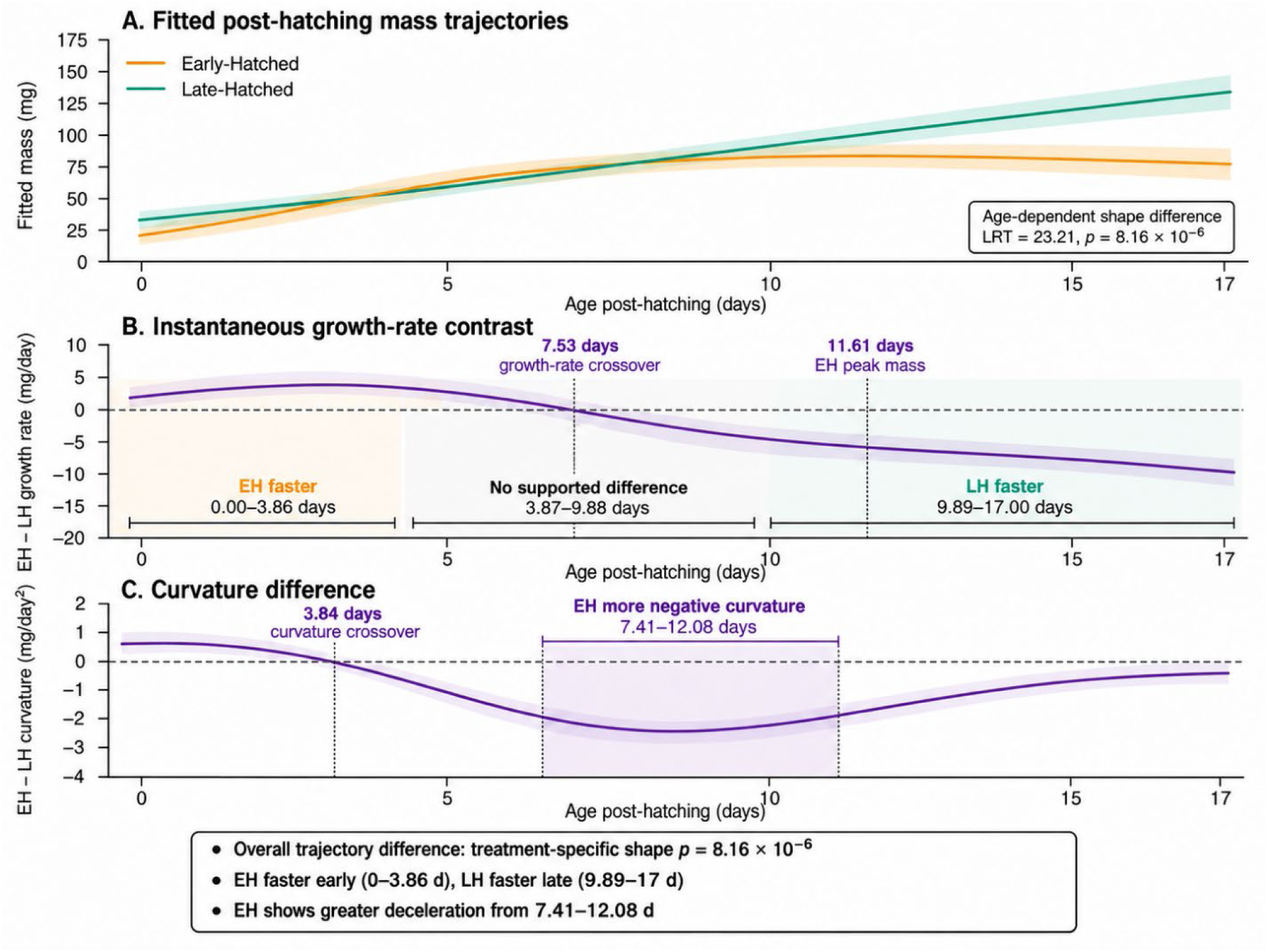
Treatment-specific post-hatching growth trajectories and derivative contrasts in Early-Hatched and Late-Hatched tadpoles. (**A**) Unanchored conditional-median mass predictions from the Gaussian log-mass GAM fitted to all 173 mass-valid EH/LH animals, with treatment-specific thin-plate smooths (k = 4), maximum-likelihood smoothing selection, and family random effects set to zero for prediction. Treatment-specific age smooths improve fit relative to a common smooth with a treatment main effect (approximate LR = 23.21, Δdf = 1.93, p = 8.16 × 10⁻⁶). Curves are displayed over shared APH support of 0–17 days; ribbons are pointwise 95% intervals. (**B**) EH−LH first-derivative contrast on the conditional-median mass scale, in mg/day. The approximate simultaneous 95% band is above zero over 0–3.86 APH, includes zero over 3.87–9.88 APH, and is below zero over 9.89–17 APH. The fitted treatment slopes cross near 7.53 APH. The EH fitted slope reaches zero near 11.61 APH, marking its mass maximum. (**C**) EH−LH second-derivative contrast, in mg/day². The simultaneous band is below zero over 7.41–12.08 APH, indicating more negative EH curvature during that interval; the point-estimate contrast crosses zero near 3.84 APH. Derivative bands use smoothing-uncertainty-corrected covariance, delta-method gradients, and 20,000 Gaussian coefficient-error simulations (seed 20261031) across a 0.01-day grid. Bands are simultaneous over age separately for each derivative order. Localized intervals depend on spline flexibility; the later LH slope advantage persists at k = 6 and k = 8, whereas the early positive contrast and negative-curvature interval do not retain simultaneous-band support.

*Second-Derivative Results* — Second-derivative contrasts showed significantly greater EH deceleration from 7.41–12.08 days APH, indicating more negative curvature of the fitted EH mass curve during this interval. The estimated curvature contrast crossed zero near 3.84 days APH [Figure 8c]. Thus, early hatching was associated with rapid initial growth followed by pronounced mid-developmental deceleration, whereas late-hatched tadpoles exhibited slower early growth followed by sustained mass gain [Figure 8c].

These localized findings were sensitive to spline flexibility, as described below.

GAM Sensitivity Analyses — Treatment-specific trajectories remained supported with k = 6 and k = 8 (approximate p = 1.05 × 10⁻⁵ and 9.59 × 10⁻⁶, respectively). These fits retained a later LH slope advantage, beginning at approximately 10.72 and 11.28 days APH, but neither identified an interval of greater EH slope or more negative EH curvature using simultaneous bands. Excluding the five-record QC batch in the k = 4 model also retained the overall trajectory difference (N = 168; approximate LR = 29.24, Δdf = 1.98, p = 4.30 × 10⁻⁷); its first-derivative bands supported EH > LH over 0–6.18 APH and LH > EH over 9.53–17 APH. Thus, overall trajectory differences and the later LH slope advantage were retained across these sensitivity analyses, whereas the early and curvature intervals depended on the specification.

### Post-Hatching Body Size and Shape

Complete morphometric data were available for 32 EH and 35 LH tadpoles. Early-Hatched tadpoles were, on average, sampled later after hatching than Late-Hatched tadpoles (*M* = 12.44 days, *SD* = 5.07, 95% CI [10.61, 14.26] vs. *M* = 9.94, *SD* = 4.73, 95% CI [8.32, 11.57]); therefore, inference was based on age- and family-adjusted models rather than unadjusted group means alone [Table 1].

**Table 1.** Body Morphology in Early and Late-Hatched *A. callidryas*. Values are unadjusted means (SD) with t-based 95% confidence intervals. Morphometric dimensions are based on 32 Early-Hatched and 35 Late-Hatched tadpoles. Harvesting mass was available for 27 Early-Hatched and 34 Late-Hatched tadpoles.

| Measure | Early-Hatched, *M* (*SD*) [95% CI] | Late-Hatched, *M* (*SD*) [95% CI] |
|---|---|---|
| **Harvesting mass (mg)** | 73.33 (17.48) [66.42, 80.25] | 85.68 (39.04) [72.05, 99.30] |
| **TTL (mm)** | 17.89 (3.31) [16.70, 19.08] | 18.24 (3.78) [16.94, 19.54] |
| **Body length (mm)** | 5.93 (1.00) [5.57, 6.29] | 6.20 (1.35) [5.74, 6.67] |
| **Body height (mm)** | 2.69 (0.56) [2.49, 2.89] | 2.90 (0.60) [2.69, 3.10] |
| **Tail-muscle height (mm)** | 1.64 (0.27) [1.55, 1.74] | 1.88 (0.40) [1.74, 2.02] |
| **Tail-fin height (mm)** | 2.94 (0.79) [2.65, 3.22] | 3.35 (0.69) [3.11, 3.59] |
| **Tail-fin:tail-muscle ratio** | 1.78 (0.32) [1.66, 1.89] | 1.81 (0.32) [1.70, 1.92] |

The family-adjusted model indicated that overall size differed by hatching treatment across post-hatch development, *F*(3, 53) = 6.92, *p* = .0005. Evidence that the groups differed specifically in the age dependence/trajectory (joint linear and quadratic interactions) was suggestive but did not meet the conventional threshold for significance, *F*(2, 53) = 2.63, *p* = .081. At a common age of 10 days post-hatch, the family-averaged predicted geometric-mean linear size was 4.31 mm (95% CI [4.01, 4.64]) for EH tadpoles and 4.56 mm (95% CI [4.28, 4.86]) for LH tadpoles. The corresponding LH/EH size ratio at APH 10 was 1.058 (95% CI [0.958, 1.169], p = .260), so the difference between these point predictions was not statistically supported at that age.

In contrast, proportional body shape did not differ significantly by hatching treatment after accounting for age, overall size, and Family of Origin (omnibus permutation F(2, 54) = 0.96, p = .449), nor was there evidence of a treatment-specific shape trajectory (F(1, 54) = 1.37, p = .246). The first three shape PCs accounted for 93.4% of the variation in size-independent morphology. None of the targeted relative body or tail dimensions, including the tail-fin: tail-muscle-height ratio, showed a significant treatment-associated profile (the smallest parametric *p* =.174).

Thus, the morphometric data indicate treatment-associated differences in overall size across development without evidence for a broad treatment-associated reorganization of proportional body shape. Explicit TTL measurements provide a complementary description of absolute linear size, whereas the multivariate analysis incorporates information distributed across TTL, body dimensions, and tail dimensions while accounting for age and family.

### Differential Gene Expression

The first two components of our PCA accounted for 34.9% and 19.9% of the total variance in gene expression across samples. Along the first axis, samples clustered by clutch, with older sibling pairs displaying higher values. The second axis was correlated with treatment, especially when considering individual sibling pairs [Figure 9a], suggesting an appreciable effect of hatching time on gene expression. That being said, the difference in PC2 between differently-treated siblings paralleled their mass at sampling [see Figure 3], which indicates that overall developmental differences may also have contributed to PC2.

**Fig. 9:**
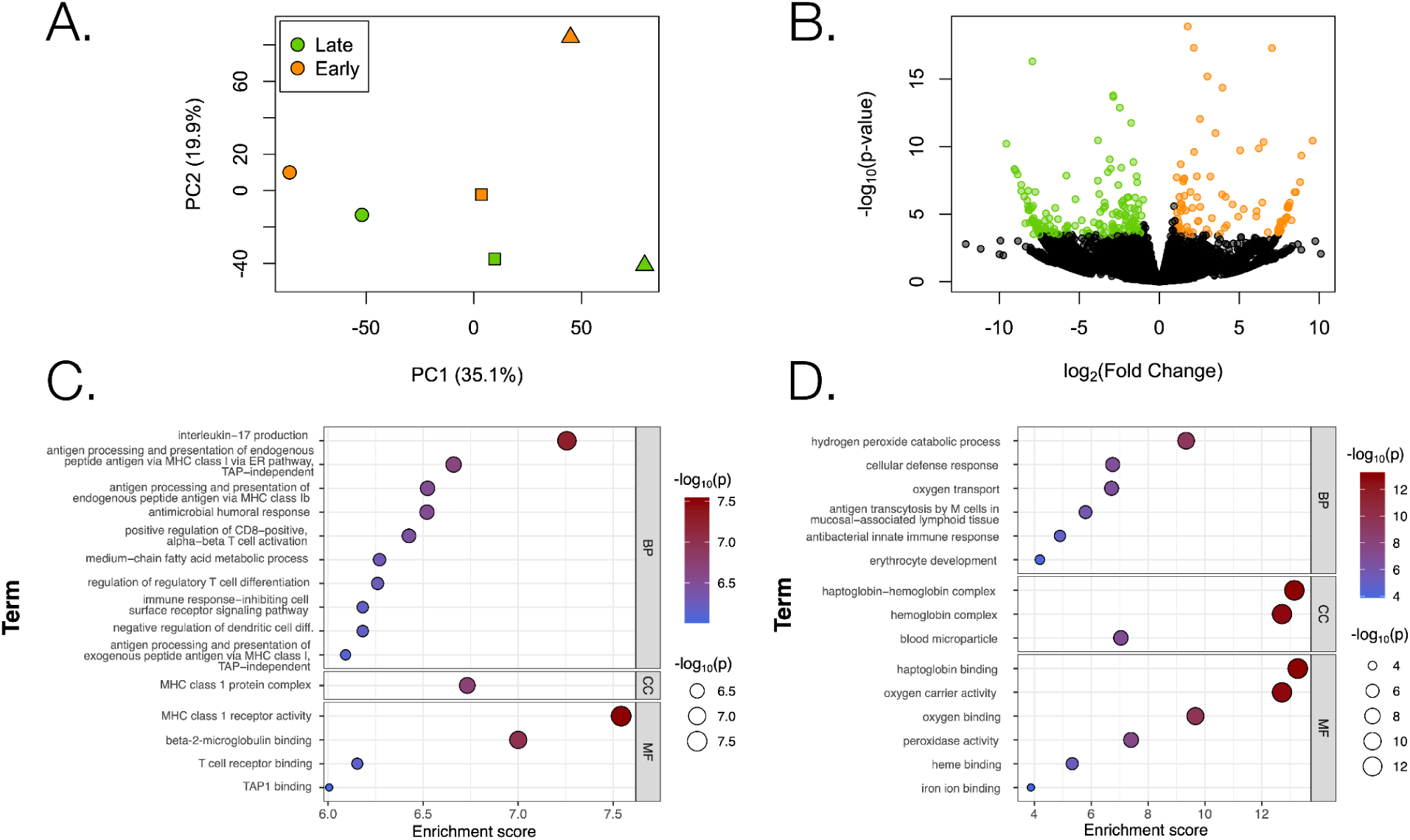
Gene expression differences associated with stress-induced early hatching. **(A)** Principal Component Analysis (PCA) based on overall gene expression shows individuals cluster by clutch (indicated by point shape, symbols correspond to those in Fig. 3), rather than treatment (NH and EH - indicated by green and orange colors respectively). **(B)** Volcano plot displays differentially expressed genes in NH tadpoles (green points) and EH tadpoles (orange points) based on a greater-than-twofold expression difference, equivalent to an absolute log₂ fold change greater than 1, and adjusted p-value cutoff of 0.05. GO term enrichment shows higher relative expression of adaptive immune response related genes in NH tadpoles **(C)** and higher relative expression of oxygen transport and oxidative stress related genes in EH tadpoles **(D)**.

Differential expression analysis yielded 258 differentially expressed transcripts (FDR-adjusted p<0.05; fold-change > 2) [Figure 9b]. Of these, 89 had higher expression in EH, and 169 in LH tadpoles. Transcripts with higher expression in late-hatching tadpoles were enriched for GO terms associated with adaptive immune function and include major histocompatibility complex related genes such as ZA2G and MR1 [Figure 9c], while those with higher expression in early-hatched tadpoles were enriched for innate defense GO terms, as well as those related to oxygen transport and H_2_O_2_ catabolism, including hemoglobin subunits and uromodulin [Figure 9d]. A list of over and under expressed genes and their corresponding GO terms is available as supplementary material (**Tables S1**).

## DISCUSSION

The pace of developmental maturation, including hatching, growth rate, and the timing of metamorphosis, is phenotypically plastic (Larsen et al., 2019). An inordinate number of interactions between genes and environmental factors (humidity, temperature, predation, exposure to wind or sunlight) force adaptation to fluctuating micro- and macroclimatic conditions. A diverse life-history portfolio supports species survival in the face of challenging social or ecological perturbations (Middleton et al., 2019). For organisms with mandatory parental care, adjustment of the parameters for developmental timing can facilitate the necessary overlap between generations.

Deciphering developmental timing patterns of growth and gene expression in a new experimental model system, the red-eyed treefrog (*Agalychnis callidryas),* may illuminate early stress-mediated accelerated early maturation across species, including humans.

This study expands our understanding of precocious development in an amphibian model of early life stress. We asked whether morphological features, such as body mass, would reflect the demands of hatching early (escape-hatching, in response to a stressor) or late (hatching undisturbed). Our data show that stress-induced EH alters the temporal organization of post-hatching development in *A. callidryas*, and results in marked physiological differences. Three findings were particularly informative and concern the growth, body morphology, and transcription response to our experimentally induced ELS. Now, each will be discussed in turn.

### I. Growth: EH and LH tadpoles followed significantly different post-hatching mass trajectories. EH tadpoles exhibited rapid early mass gain followed by pronounced deceleration, whereas LH tadpoles exhibited slower initial but more sustained mass gain

To examine the impact of stress-induced early hatching on the developmental timing of growth, harvesting weights from 173 early-hatched (EH) and late-hatched (LH) tadpoles were used in cross-sectional analyses of body mass by age since laying (APL) and age since hatching (APH).

#### Differences in Mass -

Body mass was positively associated with age in both treatments. When age was measured from laying, evidence for a treatment difference in linear slope depended on the variance estimator: the conventional test did not meet the .05 threshold, whereas HC3 inference provided modest evidence for a difference. When age was measured from hatching, the steeper LH mass–age association was supported by both approaches and persisted within the shared age range. These are population-level patterns across individuals sampled at different ages; longitudinal measurements are needed to determine whether they reflect differences in individual growth or compensatory responses.

#### Differences in Growth Dynamics -

The segmented analyses suggested that treatment differences in mass–APH associations changed with age. The first-week point estimates favored a steeper EH slope, but support depended on the covariance estimator. During days 8–14, the LH slope was clearly steeper. At ≥15 days, neither the separate-window nor continuous-piecewise analysis identified a reliable between-treatment slope difference, and late-stage estimates were imprecise. Thus, the apparent treatment contrast depended on both the age interval and the modeling approach; these results do not establish equivalent chronological-age trajectories or a uniform treatment effect throughout post-hatching development.

The timing of hatching appears to shape subsequent growth dynamics in a way that is obscured when individuals are compared only by time since laying. The improved model fit under APH further supports the interpretation that hatch timing is a meaningful developmental reference point for evaluating tadpole growth. Here, segment-based and derivative analyses showed that the treatment effect was dynamic rather than constant across development. EH tadpoles were characterized by rapid early post-hatching mass gain followed by pronounced deceleration and an eventual plateau and decline.

Conversely, LH tadpoles exhibited slower initial growth followed by sustained and increasingly greater mass gain later in the post-hatching period. Arguably, these estimates should be interpreted cautiously because of smaller sample size and shorter late-stage coverage for LH individuals. Nevertheless, the segmented analyses imply that group divergence was not constant through time; but instead reflected an active shift in growth patterns after hatching. Overall, the findings point to a time-dependent growth response. On the one hand, early hatching has a demonstrated effect of conferring initial growth advantage. This early compensatory growth during the first week post-hatching in EH likely serves to support their ability to live-on in their second niche (the pond). On the other hand, EH eventually demonstrate reduced growth potential compared to LH tadpoles, which exhibit stronger or more sustained growth in body mass.

#### Growth-Trajectory Differences -

EH and LH tadpoles differed not only in the magnitude of growth at certain ages but in the temporal organization of post-hatching growth itself. EH tadpoles followed a pronounced nonlinear, inverted-U-like trajectory: rapid early growth was followed by significant deceleration, with fitted mass reaching a maximum at approximately 11.6 days APH and declining slightly thereafter. In contrast, LH tadpoles showed a sustained monotonic increase in mass across the common observation period. Although the LH trajectory appears curvilinear when expressed in milligrams, it was approximately linear on the log-mass scale used for modeling, consistent with continued proportional growth rather than the rise-and-deceleration pattern observed in EH tadpoles. With the data being cross-sectional, these fitted patterns characterize the sampled population; they do not directly measure within-individual growth or demonstrate that EH animals lost mass over time.

The first-derivative analysis demonstrated that the relative growth advantage reversed during development. EH tadpoles grew significantly faster during approximately the first four days after hatching, whereas LH tadpoles grew significantly faster beginning at approximately day 10 APH. The second-derivative analysis localized the principal difference in trajectory shape to approximately 8–12 days APH, when EH growth underwent significantly greater deceleration than LH growth. Thus, hatching treatment was associated with a shift in the timing and shape of post-hatching growth rather than a uniform increase or decrease in the cross-sectional mass–age slope association.

These results complement the segmented growth analyses by showing that the apparent early EH advantage and later LH advantage represent components of a continuous developmental transition. The GAM and derivative analyses therefore provide evidence that early versus late hatching is associated with distinct post-hatching growth strategies: accelerated early growth followed by deceleration in EH tadpoles versus delayed but sustained growth in LH tadpoles.

### II. Morphometry: Physical differences were expressed principally in overall size rather than size-independent body shape

Previous literature suggests that body morphology may be exquisitely adapted to the environment (Touchon and Wojdak, 2014). In our study, age- and family-adjusted models supported a treatment-associated overall size profile across the sampled APH distribution. The predicted size at APH 10 was modestly greater for LH, but the contrast at that age was not statistically supported. Evidence for treatment-specific linear and quadratic age effects was also inconclusive. Thus, the significant joint size-profile test should not be interpreted as independently establishing a different size-growth tempo.

In contrast, there was little evidence that hatching treatment altered proportional morphology. After accounting for age, overall size, and family of origin, EH and LH tadpoles did not differ significantly in multivariate body shape or in individual size-adjusted body and tail dimensions. Together, these findings indicate that experimentally altered hatching timing affected the tempo of somatic growth but did not produce broad changes in body proportions. Thus, developmental plasticity associated with hatching timing appears to be expressed principally through differences in growth and overall size rather than through substantial remodeling of tadpole morphology.

### III. Gene expression: Early hatching results in differential expression of genes related to immune function, oxygen transport, and oxidative stress response

To interrogate the transcriptome response to early-versus late-hatching, RNAseq was conducted on specimens that were collected during the 12 days post-hatching, from three sibling-pairs of age-matched, differently treated LH (N=3) and EH (N=3) tadpoles. In total, 258 genes were differentially expressed during this timeframe, when EH animals showed accelerated immediate growth followed by attenuated growth compared to LH.

Most differentially expressed genes were related to immune function (broadly defined). Genes related with adaptive immune functions, such as T-cell differentiation and memory (e.g. ZA2G, HMR1) showed lower expression in EH tadpoles, while those involved in innate pathogen defense mechanisms (e.g. UROM, PRB3) had higher expression in EH tadpoles. This suggests a shift in pathogen defense strategy associated with early hatching, at least in early developmental stages. Innate pathogen defense is more prevalent at earlier developmental stages in vertebrates (Divangahi and Kaufmann, 2026; Riera Romo et al., 2016), including frogs (Paiola et al., 2023a, 2023b; Ramsay and Rohr, 2023). Hence, this result is consistent with EH tadpoles experiencing a delayed onset of innate immune function in comparison to LH, perhaps as a tradeoff to fulfill the energetic requirements of faster early growth, or other “urgent” demands that arise due to early hatching. This being said, some of the immune-related genes identified also play roles in other possibly relevant functions, such as the involvement of ZAG2 in adipose lipid metabolism (Gong et al., 2009). Our results therefore represent a first look at broad patterns revealed by GO term analysis, rather than an in-depth analysis of molecular function changes associated with early hatching.

Several genes related to oxygen transport and oxidative stress reduction, including multiple haemoglobin subunits and reelin, were also upregulated in EH tadpoles. This falls in line with our observations that EH tadpoles experience faster growth upon hatching, which would elevate the demand for oxygen across tissues in support of growth, and may in turn require increased protective measures in the cellular environment to prevent oxidative damage.

## Significance of differential gene expression results

Our results suggest that stress-induced early hatching is followed by a coordinated molecular program that may enable compensatory developmental acceleration in post-hatching tadpoles. Rather than reflecting a nonspecific increase in developmental rate, differences in transcriptional profile between treatments point to changes in specific functions associated with rapid growth, and a possible tradeoff in pathogen defense system development. These results are consistent with a scenario where compensatory acceleration following stress-induced early hatching prioritizes traits important for immediate survivorship outside the egg, while delaying the onset of processes beneficial in the longer term, such as a well-formed adaptive immune system.

As evidenced by the slower growth of EH tadpoles compared to LH after the first week post-hatching, the initial growth acceleration may come at a physiological cost. The downregulation of genes involved in xenobiotic metabolism and adaptive immunity may be indicative of a slow-down in immune system development, possibly due to resources or regulatory priority being shifted away from defense and maintenance functions during compensatory development. Despite gains through benefits in early growth, adaptive plasticity or functional maturation, this tradeoff could eventually leave EH tadpoles more vulnerable to pathogens, toxins, or injury.

## CONCLUSION

The clearest treatment effect emerged when tadpoles were aligned by time since hatching rather than time since oviposition. Chronological age post-laying partially obscured the developmental consequences of entering the aquatic environment at different times, whereas APH synchronized animals relative to this ecological transition. The GAM analysis showed that EH and LH trajectories differed significantly in shape.

EH tadpoles initially gained mass more rapidly, with a statistically supported EH advantage through 3.86 APH, but underwent marked deceleration during approximately 7.41–12.08 APH. By day 10 APH the relative advantage had reversed, and LH tadpoles showed significantly greater instantaneous mass gain through the end of the shared 17-day observation period. The segmented models produced the same qualitative pattern, although first-week inference was sensitive to the covariance estimator. Thus, the principal finding goes beyond an effect of early hatching on growth rate, to a shift in the timing of mass accumulation. This EH advantage may represent an adaptive response to premature entry into the aquatic larval niche. Earlier-hatched embryos encounter locomotor, respiratory, feeding, and predation demands sooner than siblings remaining within the egg, potentially favoring rapid acquisition of larval function. Exploratory gene expression analyses suggest that EH is accompanied by coordinated transcriptional differences in immune system development and oxygen transport.

Together, these results indicate that environmentally induced EH is associated with a broader reorganization of early larval development rather than simply a shift in the date of emergence from the egg. They show that stress-induced early hatching in *A*. *callidryas* is associated with a reorganization of early post-hatching development. EH tadpoles exhibited a transient burst of mass gain followed by greater developmental deceleration, whereas LH tadpoles subsequently showed more sustained growth.

These differences were expressed primarily in overall size rather than broad proportional body shape and were accompanied by differential gene expression patterns that suggest differential investment in different axes of organismal function. The later reversal of the growth advantage raises the possibility of developmental trade-offs following premature niche transition, although fitness costs were not directly measured.

*A. callidryas* therefore provides a promising system for experimentally linking environmentally cued hatching plasticity to subsequent growth trajectories and molecular developmental responses.

Several limitations should temper interpretation of these findings. First, each tadpole was measured only once. The mass–age relationships therefore represent population-level, cross-sectional developmental trajectories rather than longitudinal growth of individual tadpoles. The time-dependent differences between EH and LH animals demonstrate treatment-associated differences within the common experimental and husbandry environment, but they should not be interpreted as establishing a canonical growth trajectory for *A. callidryas* or as direct estimates of individual compensatory growth.

Localized derivative intervals were sensitive to spline flexibility and uncertainty estimation and should be interpreted as model-based timing estimates rather than exact developmental boundaries. In addition, tadpoles were sampled within family and clutch groups, and the primary OLS mass models did not account for within-group dependence. HC3 addresses unequal residual variance but is not a correction for family or rearing-unit clustering. The experimental unit and shared housing structure should therefore be considered when evaluating precision and generalizability.

Second, wet mass and linear morphometry provide different and complementary descriptions of larval phenotype and are subject to different sources of measurement variation. Wet mass of small aquatic larvae is particularly sensitive to hydration and residual surface water, whereas total length may be affected by specimen position, curvature, or tail condition. Comparison with previously published *A. callidryas* data indicates that, while our linear measurements overlap more closely with previously reported lengths at comparable early developmental stages, some of our early wet-mass measurements are higher than values reported under other rearing and sampling conditions. Interestingly, this published literature itself shows substantial variance later in larval development. For example, animals studied by Bouchard and colleagues (2016) reached ∼45 mm/700 mg on very different schedules depending on competition. It is fair to argue that published values are useful context, but not a canonical growth standard against which other sample populations must conform. We therefore interpret the absolute wet-mass values cautiously and emphasize comparisons between EH and LH animals maintained and sampled under the same experimental protocol.

Importantly, our independently measured morphometric dataset provides a complementary phenotype: explicit TTL measurements describe absolute linear size, while age- and family-adjusted multivariate analyses indicate treatment-associated differences in overall size without evidence for broad differences in size-independent of body shape. These qualifications do not alter the primary purpose of the study, which was to examine whether experimentally altered hatching timing was associated with coordinated phenotypic and transcriptional differences under a common laboratory environment. The observed mass–age and morphometric patterns provide phenotypic context for the transcriptomic comparison rather than an estimate of species-wide growth norms. Future studies using longitudinal measurements under varied ecological and husbandry conditions could determine the extent to which these developmental trajectories generalize beyond the present experimental setting.

Third, we did not directly measure swimming performance, predator avoidance, survival, or subsequent reproductive performance. Consequently, the ecological and functional significance of the treatment-associated growth trajectories remains unresolved. Direct measures of performance and fitness would be required to determine whether the observed phenotypic and transcriptional differences translate into ecological costs or benefits. Future studies combining longitudinal growth measurements with direct measures of locomotor performance, predator avoidance, survival, and later-life fitness will be necessary to determine whether differences in developmental timing and growth translate into meaningful ecological consequences.

Finally, given our sampling scheme and the lack of high-quality genomic resources for our study species, the transcriptomic results should be interpreted with care. RNA-seq was conducted in three EH/LH sibling pairs, and pairs were sampled across a range of developmental ages. This resulted in a strong correlation between developmental timing and global expression, which could also reflect family structure. Furthermore, because samples were not taken at individual developmental stages, what we have captured are the effects of early hatching on gene expression over the first few weeks, rather than the effector genes that trigger changes associated with stress-induced early hatching.

Finally, since functional interpretation of gene expression data hinges on an accurate annotation of transcript function and given that our transcript annotations are based on sequence similarity to proteins whose function was elucidated in distantly-related organisms (often mammals), the RNA-seq findings are best viewed as identifying candidate genes and pathways for further study in targeted functional experiments.

Despite these shortcomings, our study’s significance is two-fold. First, it elucidates the cellular-molecular mechanisms underlying life-preserving developmental trajectories. Late and Early-hatching similarly represent “niche-shifting” behavior that occur at different time points. EH has the additional feature of being prompted by some threat of predation or disturbance. EH therefore potentially highlights epigenetic factors modulating trade-offs between escape or late-hatching among red-eyed treefrogs.

Secondly, by identifying a comparable transcriptional response to threat, our study invites further interrogation of the evolutionary developmental neurobiology and ecology of stress in all living organisms. Using an ecologically valid model, we highlight potential mechanisms by which early compensatory developmental responses can lead to long-term morbidity and mortality.

Lastly, we see connections between this and human studies that recently identified a Conserved Transcriptional Response to Adversity Model (CTRA) associated with increased inflammation and accelerated aging in those experiencing despair from chronic stress of social isolation (Cacioppo et al., 2015a; Cole et al., 2015; Yang et al., 2024). Cacioppo and colleagues have argued for expanding the study of biomarkers of loneliness across phylogeny (2015b). In parallel with their human research, our current findings support a model in which stress triggers a developmentally plastic response that supports immediate survival by reallocating molecular investment away from systems that protect longevity or promote senescence (Divangahi and Kaufmann, 2026). In a sentiment shared by other researchers on the ontogeny of immunity, Romo, Pérez-Martínez and Ferrer (2016) wrote that “[t]he understanding of innate [immunological defense] mechanisms in non-mammalian vertebrates is the key to comprehend the general picture of vertebrate innate immunity and its evolutionary history” (Paiola et al., 2023b, 2023a; Ramsay and Rohr, 2023). In conclusion, *A. callidryas* represents a good model system for elucidating the epigenetic mechanisms by which ELS alters life history, with important implications for future prevention, restoration, and conservation.

## ACKNOWLEDGEMENTS

### Author Disclosures

None.

### Institutional Animal Care and Use Committee (IACUC) Approvals

This research was conducted under Trinity College IACUC Protocol Numbers: P67-22 and P67-25 as well as Marine Biological Laboratory (MBL) IACUC Protocol Numbers: 22-27, 23-32, and 24-34.

### Funding & Grant Support

SBS: Ernest Everett Just Fellowship and Whitman Fellowship, Marine Biological Laboratory (MBL); Faculty Research Committee, Trinity College; Culture, Mind & Brain Research Network, Foundation for Psychocultural Research (FPR).

## Acknowledgements

The authors would like to thank the MBL Community (Dr. Anne Sylvester, Briana Bertochi, & Daniel Johnson), the National Xenopus Resource Center (Nikko-Ideen Shaidani, James Parente, Lori Horb, & Kelsey Coppernath), Trinity College staff (Jenny Nord, Erin Mosteller, & Kathy Mallinson), Trinity College students (Kane Woods, Cass Washington, Ashton Rivera) and Mr. Jonathan P. Harwell. We thank Karen Warkentin and Maria Jose Salazar-Nicholls of Boston University for their inspiration and support. We appreciated their sharing wisdom from extensive research experience with *A. callidryas*, their donation of custom paludariums to support our breeding, and their feedback on early drafts of this manuscript. We thank Yanileth López-Tacoaman for help interpreting immune-related gene expression results. Computation for this project was carried out using resources provided by Advanced Research Computing at Virginia Tech.

## Author Contribution Statement

Conceptualization and Research Design: SBS Planning and Methodology: SBS, MH, and RM

Study Execution and Data Collection: SBS, AH, SD, EB, MR, JA, NV, AJ, MI, and MH.

Data Analysis & Interpretation: SBS, RM, KLG, AH, and SD.

Writing – Original Draft: SBS

Writing – Review & Editing: SBS, RM, and KLG. Supervision/Funding: SBS, MH, and RM.

Final Approval: SBS, RM, KLG, AH, SD, EB, MR, JA, RR, S-SP, and MH

